# Rare germline structural variants increase risk for pediatric solid tumors

**DOI:** 10.1101/2024.04.27.591484

**Authors:** Riaz Gillani, Ryan L. Collins, Jett Crowdis, Amanda Garza, Jill K. Jones, Mark Walker, Alba Sanchis-Juan, Chris Whelan, Emma Pierce-Hoffman, Michael Talkowski, Harrison Brand, Kevin Haigis, Jaclyn LoPiccolo, Saud H. AlDubayan, Alexander Gusev, Brian D. Crompton, Katie A. Janeway, Eliezer M. Van Allen

## Abstract

Pediatric solid tumors are rare malignancies that represent a leading cause of death by disease among children in developed countries. The early age-of-onset of these tumors suggests that germline genetic factors are involved, yet conventional germline testing for short coding variants in established predisposition genes only identifies pathogenic events in 10-15% of patients. Here, we examined the role of germline structural variants (SVs)—an underexplored form of germline variation—in pediatric extracranial solid tumors using germline genome sequencing of 1,766 affected children, their 943 unaffected relatives, and 6,665 adult controls. We discovered a sex-biased association between very large (>1 megabase) germline chromosomal abnormalities and a four-fold increased risk of solid tumors in male children. The overall impact of germline SVs was greatest in neuroblastoma, where we revealed burdens of ultra-rare SVs that cause loss-of-function of highly expressed, mutationally intolerant, neurodevelopmental genes, as well as noncoding SVs predicted to disrupt three-dimensional chromatin domains in neural crest-derived tissues. Collectively, our results implicate rare germline SVs as a predisposing factor to pediatric solid tumors that may guide future studies and clinical practice.

## Introduction

Extracranial pediatric solid tumors comprise approximately one-third of all new pediatric cancer diagnoses per year and are a leading cause of childhood mortality.^1^ These cancers are often aggressive and require multimodal treatment with cytotoxic chemotherapy, surgery, and radiation. Cure rates have remained modest due to a reliance on non-targeted therapies, and patients who are cured often face substantial lifelong morbidity.^2, 3^ Progressive improvements in genomics over the last two decades have greatly expanded our understanding of the somatic and germline molecular features of these rare diseases.^4–7^ In particular, the study of germline genetics—especially rare variants—has provided insight into the biological pathways responsible for the earliest stages of tumorigenesis.^8^ cancers.^5, 14–16^ Identifying missing risk factors beyond conventional SNVs/Germline genetic factors are especially important in pediatric cancers, where patients have not lived long enough to accrue substantial exposure to environmental carcinogens like tobacco smoke or UV radiation. Epidemiological studies have estimated a 4.5-fold familial relative risk for pediatric solid tumors,^10^ yet only 10-15% of pediatric cancer cases can be attributed to currently recognizable germline risk factors.^11, 12^ This paradox implies that a large portion of the heritability for pediatric cancers is still “missing” (i.e., undiscovered).^13^ However, conventional germline genetic analyses in these diseases have often focused on small subsets of variants, specifically single nucleotide variants (SNVs) and small insertions/deletions (indels) in a narrow list of genes with preexisting evidence as risk factors in other Structural variants (SVs) are an understudied family of genomic alterations that comprises any rearrangement impacting ≥50 nucleotides. By convention, SVs are classified into distinct mutational classes based on their alternative allele structures, such as deletions or duplications (collectively: copy-number variants [CNVs]), insertions, inversions, translocations, and more complex rearrangements.^17, 18^ Somatically acquired SVs have been widely implicated in oncogenesis and tumor evolution.^19, 20^ In addition, germline SVs are now recognized as critical factors in human population genetics, gene regulation, and numerous diseases. ^21, 22^ Prior focused evaluations of rare germline SVs in patients with pediatric solid cancers have revealed hints of possible germline SV risk factors,^23, 24^ but these studies have been limited by small sample sizes, a lack of matched healthy controls, and/or reliance on low-resolution genomic technologies.

We hypothesized that germline SVs might represent a novel source of disease predisposition and oncogenesis with potential clinical impact. We therefore investigated rare germline SVs in three pediatric extracranial solid tumors—Ewing sarcoma, neuroblastoma, and osteosarcoma—using germline whole-genome sequencing (WGS) from pediatric cases and ancestry-matched adult controls. We sequentially dissected five aspects of rare germline SVs in disease predisposition (**Fig. 1A**), which collectively demonstrated that rare SVs are an important component of predisposition to pediatric solid tumors and should be considered in both translational research and clinical diagnostic testing.

**Fig. 1.**
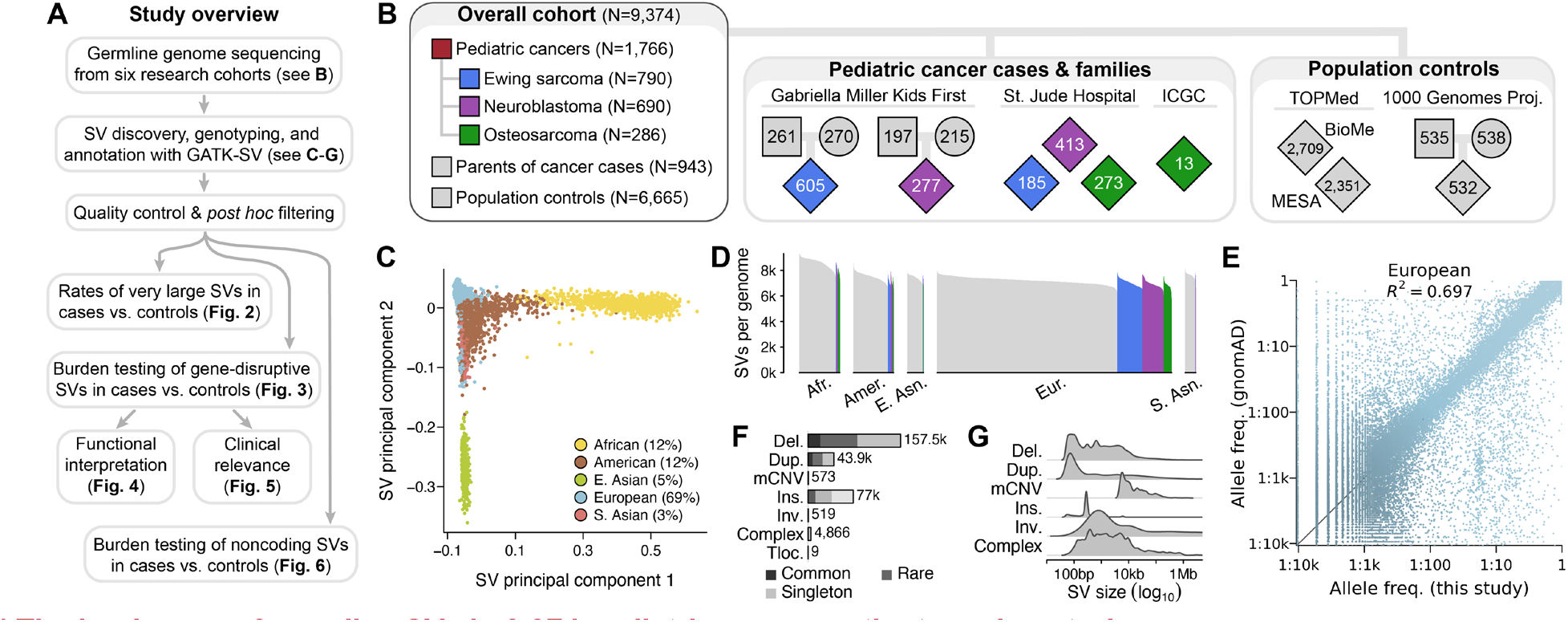
The landscape of germline SVs in 9,374 pediatric cancer patients and controls. (**A**) Roadmap of key datasets and analyses in this study. (**B**) Summary of study cohort structure after excluding low-quality WGS samples. (**C**) A principal component analysis of common, high-quality SVs stratified individuals based on their genetic ancestry. See also **Fig. S2**. (**D**) We detected an average of 7,276 high-quality SVs per genome, which was correlated with sample ancestry as expected based on human demographic history. ^18^ (**E**) The allele frequencies (AFs) of SVs detected in this study were strongly correlated with AFs of SVs reported from WGS of 63,046 unrelated individuals in the Genome Aggregation Database (gnomAD). ^17^ European samples shown here; see **Fig. S3** for other ancestries. (**F-G**) Most SVs were small (<1kb) and rare (AF<1%). Del.: deletion. Dup.: duplication. mCNV: multiallelic CNV. Ins.: insertion. Tloc.: reciprocal translocation.

## Results

### Characterization of germline SVs in 9,374 genomes

We aggregated and harmonized short-read germline WGS data (median coverage=24-fold) generated from DNA derived mostly from peripheral blood or saliva of 10,590 individuals recruited by six independent research studies (**Fig. S1**; **Table S1**). Effectively all (99%) pediatric cancer cases were sequenced by the Gabriella Miller Kids First Pediatric Research Program (GMKF) or St. Jude Children’s Research Hospital.^25–29^ We combined these cases with ancestry-matched adult controls sequenced by two cardiovascular disease studies: the Multiethnic Study of Atherosclerosis (MESA) and the Mt. Sinai BioMe Biobank (BioMe).^30^ We also included an ancestrally diverse panel of parent-child trios from the 1000 Genomes Project for ancestry inference and technical benchmarking.^31^ We performed genome-wide SV discovery, genotyping, and annotation jointly across all samples using GATK-SV, an ensemble cloud-based pipeline for population-scale WGS studies of germline SVs.^17^ Following extensive quality control, we retained a high-confidence SV dataset comprising 9,374 individuals passing all data quality filters, including 1,766 children diagnosed with one of three pediatric cancers—Ewing sarcoma (N=790), neuroblastoma (N=690), or osteosarcoma (N=286)—as well as their 943 unaffected parents and 6,665 adult population controls (**Fig. 1B**). The top six principal components derived from common SVs tracked with genetic ancestry, revealing that 31.4% of indiof samples based on their germline SV genotypes, which defined a subset of 6,728 strictly unrelated and ancestry-matched cases and controls for subsequent association testing.

We identified 284,395 germline SVs present in at least one genome, with the median individual in this study carrying 7,276 SVs (range=5,486-9,324 SVs; **Fig. 1D**). We rigorously assessed the quality of our SV dataset against multiple benchmarks, which confirmed that the properties and distributions of SVs in this study mirrored expectations set by prior non-cancer WGS initiatives (**Note S1**; **Fig. 1E**; **Fig. S3**).^17, 31^ Most SVs in our dataset (81.1%) appeared as rare alleles in the overall cohort (allele frequency [AF] <1%; **Fig. 1F**), and the average SV was relatively small (median size=281bp; **Fig. 1G**). Consistent with prior reports, the total number of SVs per genome correlated with continental ancestry, with individuals of primarily African ancestry exhibiting the greatest genetic diversity (median=8,704 SVs per genome).^18^ In summary, this dataset represented a high-confidence atlas of germline SVs from cases and ancestry-matched controls that allowed us to interrogate the role of germline SVs in predisposition of the three pediatric tumor histologies.

### Germline chromosomal dosage alterations are a sex-biased risk factor for pediatric solid tumors

We first hypothesized that large germline SVs might predispose to pediatric cancers given that large SVs are prominent risk factors for myriad other severe pediatric disorders.^32–35^ We systematically documented all large (>1Mb), rare (AF<1%) germline SVs present in the subset of 6,728 ancestry-matched cases and controls (**Fig. 2A**). We identified 143 large, rare SVs carried by 160 samples (50 cases and 110 controls; 4 samples carried two SVs each). These 143 germline SVs spanned a diverse mutational spectrum, including 71 CNVs (27 deletions and 44 duplications), 45 balanced inversions, 12 complex SVs, 10 aneuploidies (three XXY, three XYY, two monosomy X, and two trisomy 21; **Fig. 2B; Fig. S4A-B**), and five reciprocal translocations. All three pediatric cancer histologies exhibited elevated rates of unbalanced (i.e., net DNA gain or loss >1Mb) rare SVs compared to ancestry-matched controls (pan-cancer P=2.0×10^−4^; odds ratio [OR]=2.31; 95% confidence interval [CI]=1.48-3.59; N=84 SVs; logistic regression adjusted for sex, cohort, and genetic ancestry; **Fig. 2C**; **Table S2**). To ensure that this observed association was not driven by technical false positives in cases, we scrutinized the WGS evidence for all 84 unbalanced SVs, finding unambiguous support in the raw WGS depth of coverage for 97.6% (82/84) of SVs (**Fig. S5**). Moreover, 11 of the 84 SVs were carried by children whose parents were also in our final dataset, and we found that 9/11 (82%) of those large SVs were transmitted from an unaffected parent (5/9 paternal; 4/9 maternal), while the remaining 2/11 (18%) likely arose *de novo*. These *de novo* unbalanced SVs were duplications of 2.1Mb on chromosome 6 in a neuroblastoma case and 2.3Mb on chromosome 7 in a Ewing sarcoma case, neither of which overlapped any genes with established roles in cancer.^5, 14–16^ Curiously, we did not observe a similar association for the large, rare SVs that did not involve altered DNA dosage, such as balanced inversions and reciprocal translocations (pan-cancer P=0.597; OR=0.86), which suggests that the relationship between large, rare SVs and pediatric cancer predisposition is specific to gross alterations in germline copy number.

**Fig. 2.**
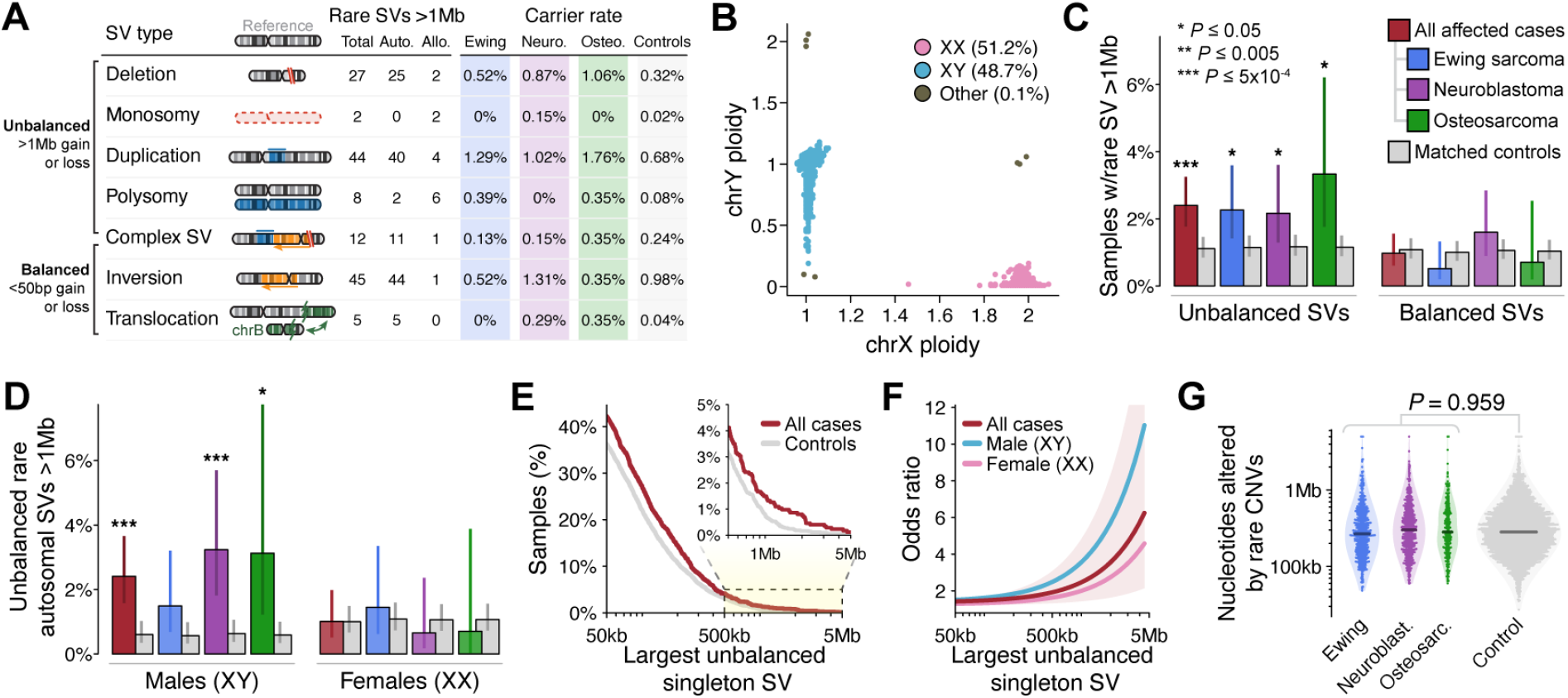
Very large germline CNVs increase risk for pediatric solid tumors in a sex-specific manner. (**A**) We identified 143 rare (AF<1%), large (>1Mb) germline SVs in a set of 1,745 pediatric cancer cases and 4,983 ancestry-matched adult controls. Auto.: autosomal. Allo.: allosomal. Neuro.: neuroblastoma. Osteo.: osteosarcoma. (**B**) WGS delineated sample sex based on ploidy (i.e., copy number) estimates of X and Y chromosomes. (**C**) We discovered an association between pediatric solid tumors and the 84 unbalanced SVs from (A). We did not observe any significant associations with comparably large but balanced rare SVs. Error bars indicate 95% confidence intervals (CI); P values derived from logistic regression adjusted for sex, cohort, and ancestry. (**D**) Results from (C) restricted to autosomes and stratified by male (karyotypic XY) vs. female (karyotypic XX) samples. (**E**) Proportion of cases and controls who carried at least one singleton unbalanced germline SV larger than the size specified on the X-axis. See also **Fig. S4F**. (**F**) Relationship between the size of unbalanced singleton SV size and the corresponding odds ratio for pediatric cancer based on the data from (D) presented as a cubic smoothing spline. Shaded area is 95% CI for a pooled model of both sexes from all histologies. (**G**) The total sum of nucleotides altered by autosomal, rare, unbalanced SVs did not significantly differ between cases & controls.

Further dissection of all large, unbalanced SVs revealed that the observed excess in cases was almost entirely concentrated in males (karyotypic XY only; pan-cancer P=9.8×10^−5^; OR=3.65), with no significant enrichment observed in females (karyotypic XX only; pan-cancer P=0.49; OR=1.27; **Fig. S4C**). This difference in effect sizes between sexes was significant (P=0.028; Welch’s t-test), and the rates of these SVs did not significantly differ between male and female controls (25/2,245 male controls vs. 29/2,472 female controls; P=0.939). These sex biases were most pronounced in neuroblastoma and persisted even after (i) excluding all SVs on sex chromosomes (**Fig. 2D**), (ii) restricting to individuals of European ancestry (**Fig. S4D**), and (iii) analyzing independent cohorts of cases separately (**Fig. S4E**). We next assessed whether the enrichment of large, unbalanced SVs in cases vs. controls was also present for smaller unbalanced SVs. The elevated rates of rare, unbalanced SVs in cases extended to relatively smaller SVs, albeit with dramatically weaker effects (e.g., pan-cancer OR=1.37 vs. OR=2.50 for singleton SVs >50kb vs. >1Mb in both sexes, respectively; **Fig. 2E-F; Fig. S4F**). Given that the median size of all unbalanced SVs in our dataset was 523bp, we tested whether these findings were simply proxies for a general genome-wide excess of rare, unbalanced SVs of all sizes in cases, but we found no significant difference in the sum of all nucleotides gained and lost due to all rare SVs between cases and controls (**Fig. 2G; Fig. S4G**). Collectively, these results established that children—especially males—are at substantially greater risk for pediatric solid cancers if their genomes harbor extensive DNA dosage imbalance due to a single large germline SV.

We reasoned that this enrichment of large, unbalanced SVs in cases could either be driven by (i) focal excesses of SVs concentrated at specific pathogenic loci or (ii) a nonspecific excess of SVs distributed uniformly throughout the genome; the former is observed in many developmental disorders,^35, 36^ whereas the latter that might be indicative of mechanisms beyond direct gene disruption, such as mitotic chromosomal stability. To distinguish between these possibilities, we first excluded all SVs that overlapped any of 621 genes with a previously established role in cancer,^5, 14–16^ finding that the original enrichment in cases remained largely undiminished (pan-cancer P=0.007; OR=2.10; N=62 autosomal SVs; **Fig. S4H**). Second, we performed case-control burden tests for all rare, unbalanced SVs ≥100kb in 1Mb sliding windows across all autosomes, which did not identify any loci surpassing Bonferroni-corrected significance (P<1.9×10^−5^) (**Fig. S6**). The lack of overlap between obvious pathogenic loci and large, unbalanced SVs in cases suggests that these SVs may impart disease risk through undiscovered pathogenic genes and/or extragenic mechanisms, like an effect on somatic genome stability.

### Gene-disruptive germline SVs are enriched in pediatric cancer patients

We next turned our attention to a different subset of SVs: focal gene-disruptive SVs, which on average were ∼250 times smaller (median size=8.0kb) but ∼60 times more abundant (N=8,638 SVs) than the large, rare SVs that were the focus of our earlier analyses. We specifically hypothesized that gene-disruptive germline SVs might also mediate risk for pediatric solid tumors. We found that cases across all three histologies had an average of 6-10 more autosomal genes disrupted by germline SVs than controls after accounting for sex, ancestry, and cohort (pan-cancer case mean=127.5 genes; control mean=120.9 genes; P<10^−10^; **Fig. 3A**). This result was not explained by outsized contributions from common SVs or large, multi-gene SVs: cases carried a significantly greater total number of rare gene-disruptive SVs (pan-cancer P=1.7×10^−5^; OR=1.07), which remained significant even when restricting to small SVs impacting single genes (P=0.002; OR=1.06; **Fig. S7A**). In contrast to the sex-biased effects of large, unbalanced SVs, we did not observe a significant difference in effect sizes of all rare gene-disruptive SVs between males and females (pan-cancer male OR=1.09; female OR=1.05; P=0.27, Welch’s t-test; **Fig. S7B**). The enrichment for gene-disruptive SVs in cases was twice as strong when restricted to singleton SVs (**Fig. 3B**) and exhibited comparable effect sizes in samples from GMKF and St. Jude cohorts when analyzed separately (**Fig. S7C**). We also noted independent enrichments of two distinct subsets of SVs predicted to have opposite consequences on gene function (**Fig. 3C**): loss-of-function (LoF), usually caused by exonic deletion (pan-cancer P=1.8×10^−7^; OR=1.19), and whole-gene copy-gain (CG) caused by duplication SVs (pan-cancer P=0.001; OR=1.32), but did not find that singleton LoF and CG SVs were mutually exclusive within individual genomes (P=0.55, Chisquare test). Taken together, these results demonstrated that the genomes of children with pediatric solid tumors tend to harbor more gene-inactivating (LoF) and gene-activating (CG) rare germline SVs than the average cancer-free adult.

**Fig. 3.**
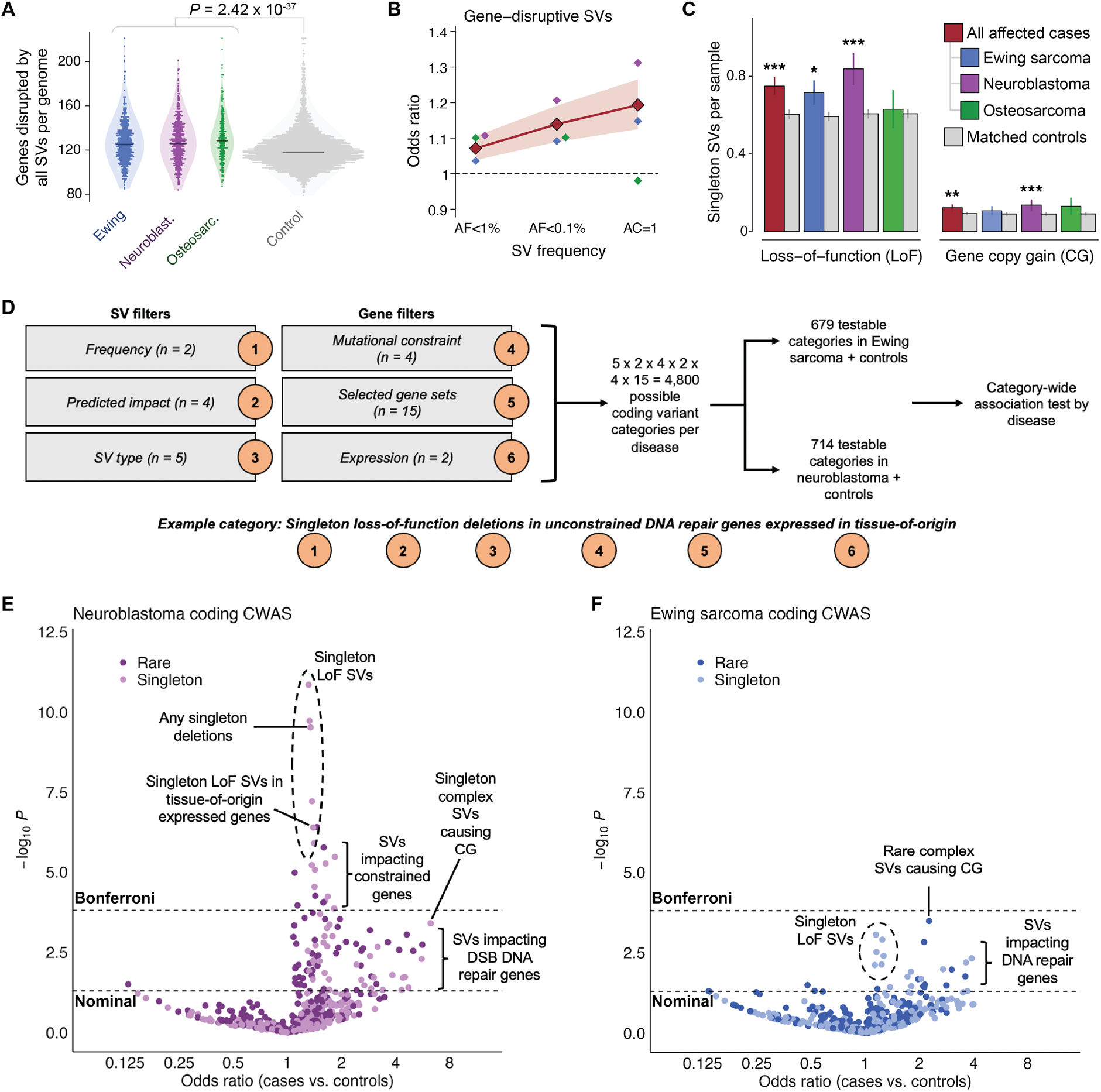
Pediatric cancer patients carry an excess of rare germline SVs that impact disease-relevant genes. (**A**) On average, the number of protein-coding genes disrupted by all germline SVs was significantly greater among pediatric cancer cases relative to adult controls. (**B**) Rare gene-disruptive SVs were enriched in cases versus controls, and this enrichment was inversely correlated with SV frequency. Shaded area indicates 95% CI for all histologies. (**C**) We found comparable enrichments in two independent subsets of singleton SVs with opposing predicted consequences on gene function: loss-of-function (LoF) and whole-gene copy gain (CG). (**D**) We carried out a category-wide association study (CWAS) in neuroblastoma and Ewing sarcoma, combining six layers of filters to categorize types of coding SVs for burden testing. ^37, 38^ We evaluated 679 and 714 categories of gene-disruptive germline SVs in Ewing sarcoma and neuroblastoma, respectively. (**E**) In neuroblastoma, 27 categories of SVs exceeded Bonferroni significance, including singleton LoF SVs impacting mutationally constrained genes and genes expressed in adult adrenal gland. (**F**) In Ewing sarcoma, no single category of SVs was enriched in cases or controls at Bonferroni significance, although multiple categories of potential biological interest were nominally significant.

To gain insight into the genes driving this global enrichment of rare LoF and CG SVs in pediatric cancer patients, we next implemented a systematic “category-wide association study” (CWAS) approach for Ewing sarcoma and neuroblastoma,^37, 38^ the two more highly powered (N≥690) histologies in our study. Briefly, we enumerated all possible combinations of six filters for our SV dataset, including filters for AF, predicted coding consequence, SV type, genic mutational constraint, gene set membership, and gene expression in the adult tissue from the Genotype-Tissue Expression (GTEx) project that was the most proximal to the putative cancer tissue-of-origin (adrenal gland for neuroblastoma and skeletal muscle for Ewing sarcoma; **Tables S3-S5**).^39^ This strategy resulted in 4,800 possible categories of SVs per disease. To focus on categories with sufficiently dense SV data, we required >10 SVs per category when summed across cases and controls; this yielded 714 and 679 “testable” categories in neuroblastoma and Ewing sarcoma, respectively (**Fig. 3D**). Within each category, we evaluated the burden of autosomal SVs in cases relative to controls after adjusting for sex, cohort, and genetic ancestry, and identified categories meeting a Bonferroni significance threshold (P≤1.6×10^−4^) after estimating the number of effective tests using permutation to account for correlation between similar categories, as has been previously proposed.^37, 38^

In neuroblastoma, we identified 27 categories of SVs surpassing our conservative Bonferroni significance threshold. Most significant categories involved LoF SVs associated with increased disease risk (e.g., singleton LoF deletions, P=3.0×10^−10^; case vs. control OR=1.34; **Fig. 3E**; **Table S6**). Among all significant categories, we observed the strongest effects for SVs causing LoF of mutationally constrained genes (e.g., singleton LoF deletions of constrained genes; P=3.4×10^−6^; OR=1.84; N=68 SVs), which intriguingly mirrors patterns reported in pediatric neurodevelopmental disorders.^40, 41^ The multi-layer design of the CWAS allowed us to further dissect these signals and identify subsets of SVs driving these broad enrichments with comparatively greater effect sizes, albeit at reduced power. We identified several smaller categories that exhibited especially strong effect sizes, such as singleton complex SVs resulting in CG of one or more protein-coding genes (P=4.0×10^−4^; OR=6.27; N=7 SVs), rare duplications resulting in CG of COSMIC cancer genes (P=0.0024; OR=2.64; N=13 SVs), and rare deletions resulting in LoF of DNA double-strand break repair genes (P=6.5×10^−4^; OR=2.88; N=14 SVs), all of which may provide insights into the types of germline SVs that confer relatively greater risk for neuroblastoma. By contrast, we found no categories surpassing Bonferroni significance in Ewing sarcoma (**Fig. 3F**; **Table S7**), although a handful of categories were nominally significant, such as singleton deletions resulting in LoF of DNA damage repair genes (P=0.0047; OR=3.91; N=8 SVs) and rare complex SVs resulting in CG of one or more genes (P=3.3×10^−4^; OR=2.26; N=22 SVs). In aggregate, this systematic CWAS of coding SVs in neuroblastoma and Ewing sarcoma supported our hypothesis that specific subsets of gene-disruptive rare germline SVs predispose to these diseases and suggested that rare germline SVs play a relatively larger role in the pathogenesis of neuroblastoma than Ewing sarcoma.

### Ultra-rare germline SVs disrupt functionally relevant genes and pathways in pediatric tumors

Given that singleton LoF SVs were the most significant risk factors identified by the CWAS in both neuroblastoma and Ewing sarcoma, we next sought to clarify their functional impact in disease pathogenesis. Particularly in neuroblastoma, sequentially filtering singleton SVs to isolate LoF deletions in constrained, tissue-expressed genes increased SV effect sizes (from OR=1.31 to 1.82 in neuroblastoma; from OR=1.15 to 1.25 in Ewing sarcoma; **Fig. 4A**). This finding reflected the guided tradeoff between power and interpretability in the CWAS framework but also implied that specific subsets of singleton SVs increased disease risk more than others. To understand biological processes that might be impacted by these genic singleton SVs, we conducted gene set enrichment analyses to identify gene ontology (GO) terms that were disrupted by singleton SVs more than would be expected by chance (N=2,722 gene sets tested). In neuroblastoma, which arises from the sympathetic nervous system,^42^ we found that gene sets involved in neurogenesis and synaptic transmission were significantly enriched for singleton gene-disruptive SVs (e.g., CNS development [GO:0007417]; OR=3.1; P=2.3×10^−6^; Fisher’s exact test; **Fig. 4B**). While less striking, we also identified several gene sets enriched for singleton gene-disruptive SVs in Ewing sarcoma that could be linked to its mesenchymal origin,^43^ including calcium homeostasis and peptide synthesis (e.g., peptide biosynthetic process [GO:0043043]; OR=4.5; P=8×10^−7^). For both diseases, we did not observe as strong an enrichment for SVs found in controls in these gene sets, suggesting that disease-specific factors mediate their enrichment rather than just genomic factors like gene size. Furthermore, we found that more specific categories with relatively greater effects also exhibited enrichment for these gene sets, highlighting that the subsets of SVs that most strongly increase risk likely also affect these pathways (**Fig. 4C; Fig. S8A-B**).

**Fig. 4.**
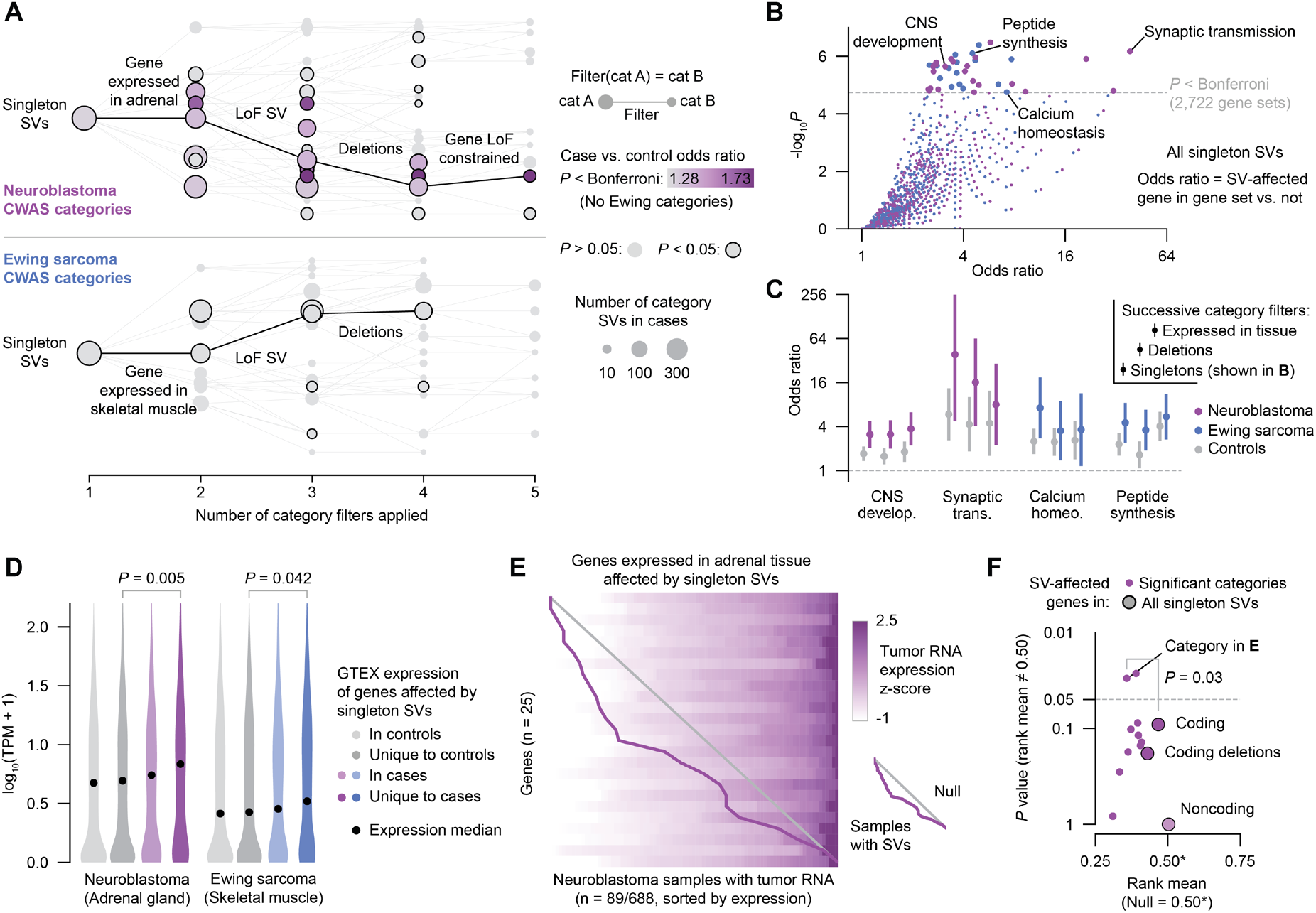
Ultra-rare germline SVs in pediatric cancer patients dysregulate gene expression in premalignant tissues and in tumors. (**A**) Diagrams of CWAS category relationships. Singleton SV categories are depicted as nodes and filters as edges. The x-axis represents the number of filters applied to each category, while the vertical space between categories is proportional to their similarity (Jaccard index). Example filter paths containing at least nominally significant categories are shown in black. (**B**) Gene set enrichment results for risk-carrying singleton SVs and GO biological processes. Odds ratios were computed by a Fisher’s exact test of an SV-affected gene being in the category and also in the gene set. (**C**) Gene set enrichment results for select gene sets and categories in cases and controls. The three categories shown for each gene set represent singleton SVs, singleton deletions, and singleton deletions affecting genes expressed in the tissue-of-origin (i.e., successively applied filters). Bars represent 95% CI. (**D**) Expression in GTEx v8 for genes affected by singleton SVs in controls, unique to controls, in cases, and unique to cases. ^39^ P values from Wilcoxon test. (**E**) Expression heatmap of genes expressed in adrenal tissue that were affected by singleton SVs in the subset of neuroblastoma patients with tumor RNA data available. Each row represents a gene and each column represents a sample; samples within each row are ordered by expression z-score. Samples with SVs (one per row) are connected by a purple line. Diagonal gray line indicates expectation under a uniform null. (**F**) Effect on tumor expression of SVs in significant neuroblastoma categories. Y-axis represents the P value of comparing to the null uniform distribution (i.e., no effect on expression). Larger points represent higher-level groupings of SVs. Indicated P value is from bootstrap-comparing the category from (E) to the background of all coding SVs.

We next reasoned that singleton germline SVs may contribute to disease risk by directly altering gene expression in the tissue-of-origin for each histology. For both neuroblastoma and Ewing sarcoma, we found that the genes disrupted by singleton SVs exclusively in affected cases were expressed at higher levels in normal tissue than those uniquely disrupted in controls, suggesting that risk-carrying SVs preferentially impact more highly expressed genes (e.g., neuroblastoma case unique vs. control unique median TPMs=6.8 vs. 4.9; Wilcoxon P=0.005; **Fig. 4D**). Most genes disrupted exclusively by singleton SVs in cases were also broadly expressed in other tissues, mirroring observations of pathogenic germline variants in severe developmental disorders (**Fig. S8C**).^41^ While RNA from normal tissue was not available from cases, a subset of neuroblastoma tumors from GMKF patients (N=89/688) had previously undergone transcriptome profiling, allowing us to assess the effect of SVs on tumor expression.^28^ To assess the overall impact of singleton SVs on genes with varied degrees of expression, we converted gene expression values to a sample-normalized percentile rank (0-1) across all 89 samples and assessed the average rank of samples with SVs. We found that singleton SVs tended to decrease expression of their affected genes in the patients’ corresponding tumor, a result that was significant for two categories (e.g., singleton SVs affecting genes expressed in adrenal tissue rank mean=0.35 vs. 0.50 uniform null, P=0.03, **Fig. 4E, Fig. S8D-E**). We also found that singleton SVs that affected genes expressed in adrenal tissue significantly decreased expression in tumors more than coding SVs generally (rank mean=0.35 vs. 0.47; permutation test P=0.03). Noncoding SVs, in contrast, did not exhibit an effect on expression of their closest gene (rank mean=0.50; P∼1; **Fig. 4F**). Taken together, these results showed that singleton SVs contribute significant disease risk and that these SVs affect genes expressed in disease tissue-of-origin, dysregulate relevant biological processes, and may even impact eventual tumor gene expression.

### Prevalence of germline SVs that disrupt cancer genes

A growing body of literature has called for WGS to become the first-tier diagnostic screen in diseases with suspected genetic etiologies.^44–46^ The clinical utility of WGS will hinge on its ability to detect pathogenic variants that are inaccessible to current technologies used in clinical diagnostics. Since germline SVs are not routinely examined in genetic testing for pediatric solid tumors and their prevalence has not been systematically examined at the resolution of WGS, we sought to understand the spectrum of SVs in our dataset that impacted recognized germline cancer predisposition genes (CPGs) and COSMIC cancer genes.^5, 14–16^ We identified 69 patients (3.5%) who carried SVs with AF<0.1% that impacted genes previously implicated in cancer predisposition and oncogenesis (**Fig. S9**), such as LoF SVs in DNA damage repair genes like *PALB2* and *BARD1* (**Fig. 5A**), which mirrors previous reports of pathogenic short variants in these same genes.^47, 48^ We additionally found that several of these SVs impacting established CPGs, like *PHOX2B* and *FANCA*,^47, 49^ were inherited from unaffected parents and therefore likely represent incompletely penetrant risk factors (**Fig. 5B**). We also noted duplications and complex SVs converging on key members of oncogenic pathways in pediatric solid tumors, such as RAS-MAPK (**Fig. 5C**),^50^ which correlated with altered gene expression in cases where tumor RNA was available (**Fig. 5D**). A subset of the rare SVs impacting CPG or COSMIC genes were low-frequency polymorphisms in the general population, including duplications of *RAF1, CHEK2, BRCA1*, and *ERCC2*, all of which were observed at greater frequencies in cases than in controls (*e*.*g*., *ERCC2* duplication: 5/773 Ewing sarcoma cases vs. 8/4,574 controls; P=0.051; OR=3.35; **Fig. 5E**).

**Fig. 5.**
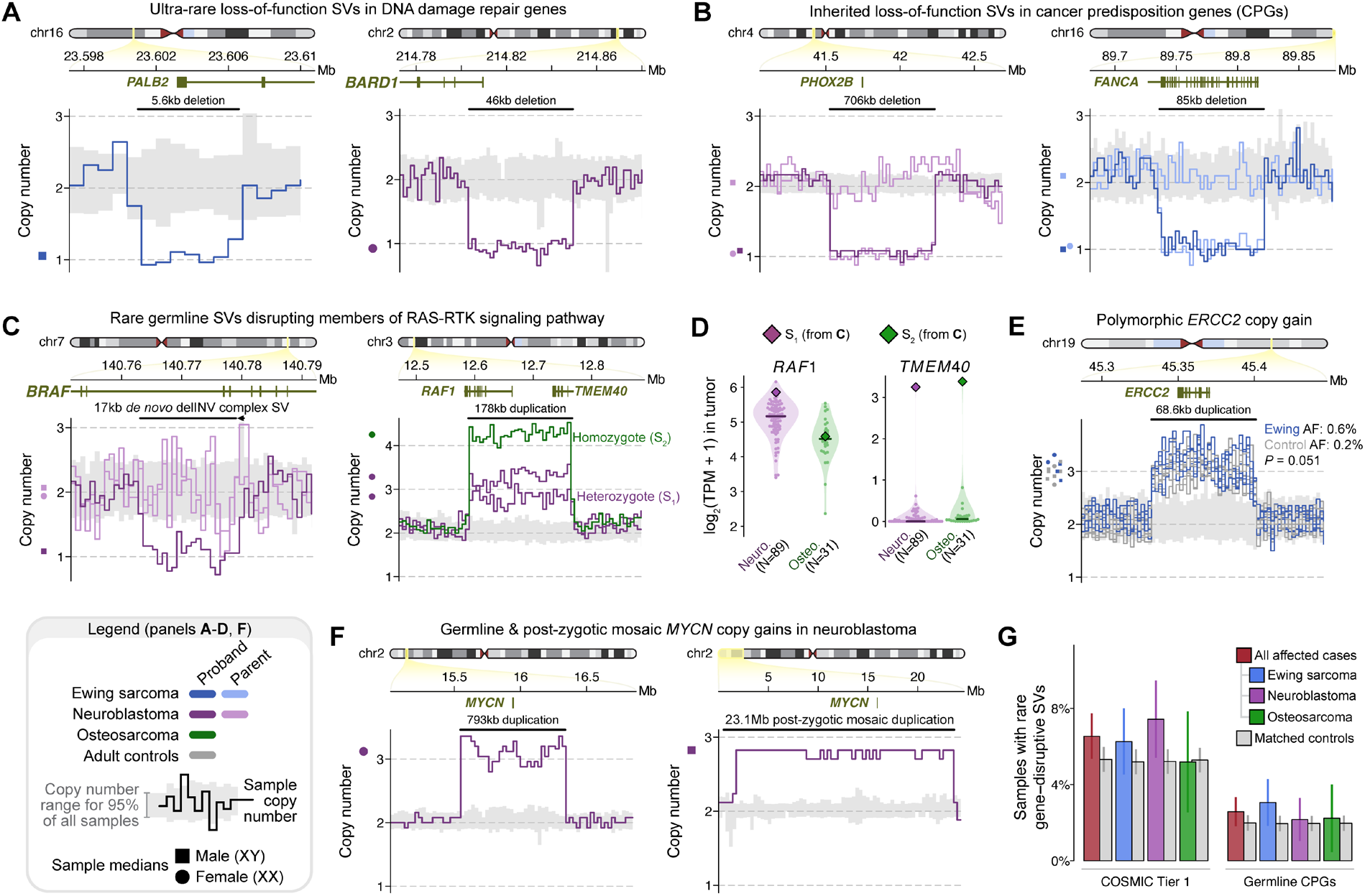
Gene-disruptive germline SVs in COSMIC and cancer predisposition genes (CPGs) in pediatric patients with solid tumors. (**A**) We found germline LoF deletions in DNA damage repair genes, such as *PALB2* in Ewing sarcoma and *BARD1* in neuroblastoma. (**B**) We also observed germline LoF SVs of known CPGs, like *PHOX2B* in neuroblastoma and *FANCA* in Ewing sarcoma, that were carried by affected children but were inherited from unaffected parents. (**C**) RAS-MAPK genes were impacted by germline SVs, including a singleton *de novo* complex SV resulting in a two-exon deletion of *BRAF* in one neuroblastoma case and an ultra-rare polymorphic duplication over *RAF1* in three unrelated cases. (**D**) The germline *RAF1* duplication from (C) was associated with high *RAF1* expression in a neuroblastoma tumor, and increased expression of *TMEM40*, the gene upstream of the *RAF1* promoter, in neuroblastoma and osteosarcoma tumors. (**E**) A rare polymorphic duplication predicted to result in CG of *ERCC2*, a DNA damage repair gene, was found at a three-fold higher frequency in Ewing sarcoma cases than in controls. (**F**) We discovered a *de novo* germline *MYCN* duplication and a likely post-zygotic *MYCN* duplication in two unrelated neuroblastoma cases. (**G**) The overall rates of gene-disruptive rare SVs in CPGs and COSMIC cancer genes were not significantly higher in cases relative to controls.

Many pediatric tumors are associated with recurrent somatic genome rearrangements, like *EWSR1-FLI1* fusions in Ewing sarcoma and *MYCN* amplifications in high-risk neuroblastoma.^43, 51, 52^ We asked whether rare germline SVs might act as precursor lesions to these recurrent somatic rearrangements, but we found no evidence of germline SVs enriched in cases versus controls at these loci (**Fig. S10**). However, we did identify a single neuroblastoma patient carrying an apparently *de novo* 793kb duplication resulting in CG of *MYCN* that was statistically consistent with a full germline event (estimated copy number=3.09; 95% CI=2.78-3.41; **Fig. 5F**). Joint analysis with the patient’s matched tumor demonstrated a high amplification (>45 copies) of *MYCN* in the tumor and no detectable tumor-in-normal contamination (**Fig. S7D**). While scrutinizing the *MYCN* locus, we also uncovered an apparently mosaic 23.2Mb CG duplication of *MYCN* in a second, unrelated neuroblastoma patient; this very large event was initially excluded by our GATK-SV pipeline due to being statistically inconsistent with a true germline event (estimated copy number: 2.79; 95% CI: 2.60-2.98), but the near-integer copy number might imply an early post-zygotic mutational origin since tumor-in-normal contamination was unlikely in this saliva-derived normal sample. While we were unable to molecularly validate the presence of these *MYCN* duplications due to the lack of additional germline DNA, we note that one other ultra-rare *MYCN* germline duplication had been reported in a neuroblastoma patient in a prior study,^23^ supporting the relevance of germline or early post-zygotic mosaic *MYCN* duplications in neuroblastoma pathogenesis.

Given the numerous examples of rare SVs predicted to disrupt established cancer genes in affected cases, we anticipated that the collective rates of rare SVs in CPGs and COSMIC genes would be elevated in cases compared to controls. However, we did not find this to be the case (CPGs, pan-cancer OR=1.16; P=0.45; COSMIC cancer genes, pan-cancer OR=1.21; P=0.12; **Fig. 5G**; **Fig. S7E**), which may imply that the totality of germline SVs across all currently recognized CPGs and COSMIC genes will not collectively explain a large fraction of the missing heritability of these pediatric solid tumors. Taken together with the enrichment of rare gene-disruptive SVs in constrained genes beyond CPGs and COSMIC genes (**Fig. S7F**), we concluded that there are likely undiscovered CPGs that confer disease risk specifically for these tumors. We systematically searched for candidate novel CPGs by conducting gene-based association tests for 15,544 genes, identifying just one candidate exceeding exome-wide significance (P<3.2×10^−6^; **Fig. S11**): rare LoF deletions impacting the promoter and first exon of *KL*, a gene previously implicated in aging that is silenced in some adult cancers,^53, 54^ were associated with sharply increased risk for neuroblastoma (P=2.2×10^−8^; OR=123.5; 95% CI lower bound=5.9). However, this candidate CPG should be interpreted cautiously in the absence of independent replication in an external dataset, and larger sample sizes will be needed to discover novel CPGs in these rare histologies.

### The contribution of noncoding germline SVs to pediatric solid tumor predisposition

Our analyses thus far had established that very large and gene-disruptive rare germline SVs were significantly enriched in pediatric cancer patients compared to controls. However, we reasoned that yet more of the missing heritability of pediatric solid tumors may be hidden in the ∼98% of the genome that does not encode proteins. To test this hypothesis, we adapted the CWAS framework to focus exclusively on SVs with no predicted coding consequence and added a functional filter capturing epigenetic and other regulatory genomic annotations (**Fig. S12N**; **Tables S8-S10**), allowing us to systematically categorize noncoding SVs in each histology. We enumerated 26,400 possible categories and found that 7,909 categories in neuroblastoma and 6,244 categories in Ewing sarcoma met our minimum “testable” threshold of >10 SVs in total across cases and controls. We once again evaluated the burden of autosomal SVs in cases relative to controls for all categories and identified noncoding categories meeting Bonferroni significance (P≤5.4×10^−5^) after estimating the number of effective tests per disease with the same permutation strategy as in our earlier coding CWAS.^37, 38^

We discovered multiple Bonferroni-significant categories associated with increased neuroblastoma risk, all of which involved noncoding singleton SVs overlapping topologically associating domain (TAD) boundaries defined from healthy adult adrenal gland tissue (**Fig. 6A**; **Table S11**).^55^ In contrast, no categories reached Bonferroni significance in Ewing sarcoma despite our CWAS test statistics being well-calibrated (**Fig. 6B; Fig. 6C**; **Table S12**). In neuroblastoma, we observed a significantly stronger effect size for noncoding singleton SVs intersecting TAD boundaries relative to all noncoding singletons (P=3.6×10^−4^, Welch’s t-test), whereas this difference was not seen in Ewing sarcoma (P=0.42) (**Fig. 6D**). Taken together, the enrichment of noncoding germline SVs overlapping tissue-specific TAD boundaries supported the potential disruption of these domains—and presumably the dysregulation of the genes residing within these TADs—as a germline risk factor for the development of neuroblastoma.

**Fig. 6.**
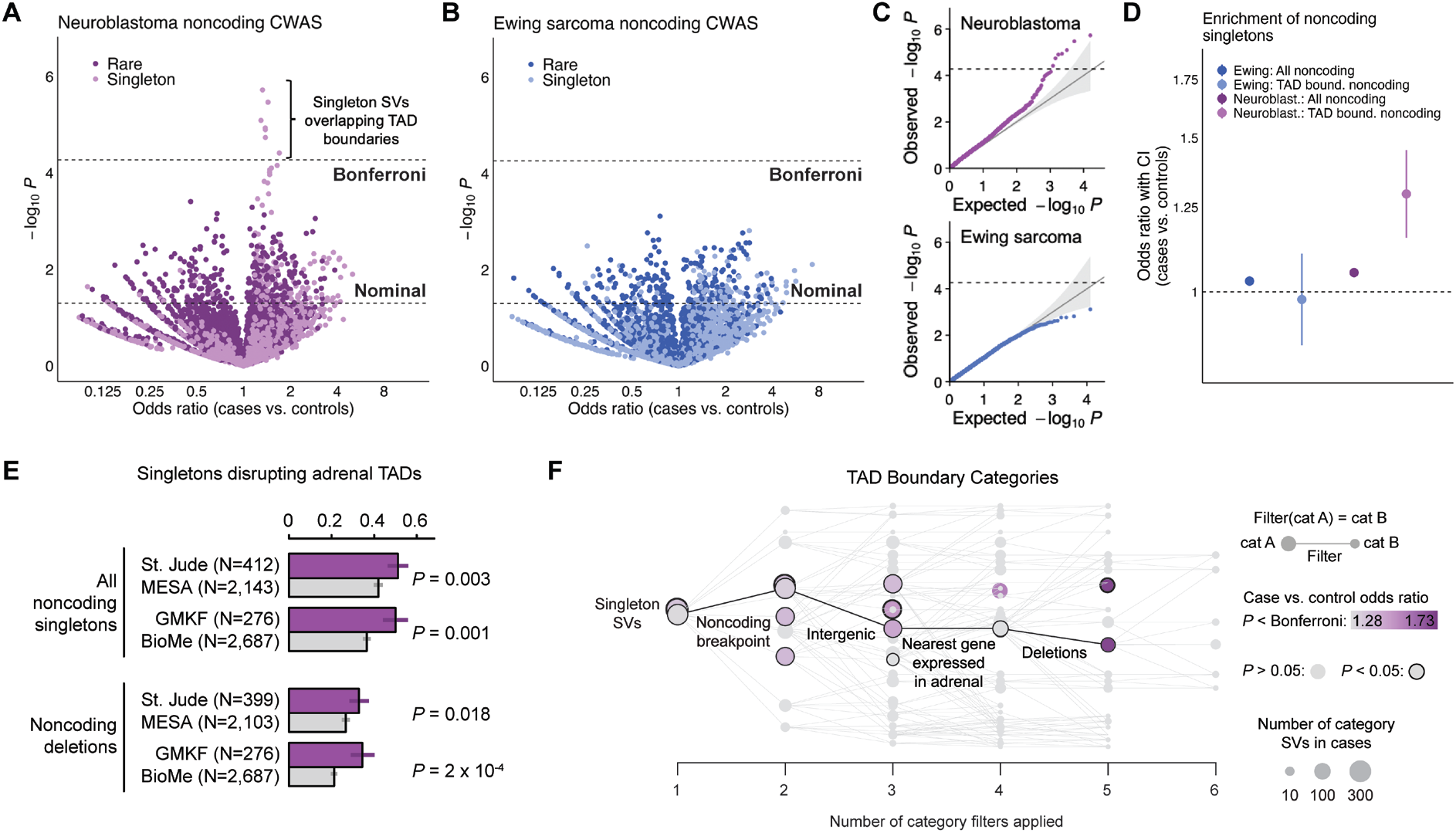
Noncoding germline SVs impacting TAD boundaries in disease-relevant tissues increase risk for neuroblastoma. (**A**)We performed a CWAS for rare, noncoding SVs in neuroblastoma cases vs. controls, finding that singleton germline SVs overlapping adrenal gland-derived TAD boundaries were significantly enriched in cases after correcting for the estimated number of effective CWAS tests (“Bonferroni”). No noncoding SV categories reached the threshold of Bonferroni significance for enrichment in Ewing sarcoma cases relative to controls. (**C**) Quantile-quantile plots for all noncoding CWAS categories for neuroblastoma (top) and Ewing sarcoma (bottom) demonstrated good calibration of CWAS test statistics. Dashed horizontal line corresponds to Bonferroni significance. (**D**) We observed a significantly stronger effect size for noncoding singleton SVs intersecting TAD boundaries defined in the putative tissue-of-origin (adrenal gland) relative to all noncoding singletons in neuroblastoma (P=3.6×10^−4^, Welch’s t-test), whereas this difference was not seen in Ewing sarcoma (P=0.42), consistent with the lack of a TAD-related association in the Ewing sarcoma CWAS. (**E**) The enrichment in neuroblastoma cases for singleton germline SVs intersecting TAD boundaries was significant in the St. Jude and GMKF cohorts when analyzed separately. (**F**) Effect sizes for singleton SVs intersecting TAD boundaries in neuroblastoma increased monotonically when subsetting to deletions in proximity to genes expressed in adrenal tissue.

We further scrutinized the noncoding TAD boundary SV categories in neuroblastoma and found that their effects were consistent in both St. Jude (P=0.003; OR=1.25) and GMKF (P=0.001; OR=1.34) when analyzed separately (**Fig. 6E**). We examined which noncoding filters resulted in the categories that carried the strongest disease risk, finding that deletions of TAD boundaries near adrenally expressed genes led to the greatest increase in SV effect sizes (**Fig. 6F**). However, we did not find any GO terms with genes that were significantly overrepresented near noncoding SVs overlapping TAD boundaries, nor did these SVs have an appreciable effect on nearby gene expression at our sample size, reflecting the major challenges of predicting the functional consequences of noncoding germline SVs on gene regulation mediated by alterations to three-dimensional chromatin domains.^56–59^

## Discussion

In this study, we systematically dissected the wide-ranging influences of rare germline SVs on pediatric solid tumor disease risk. Prior studies have implicated rare SNVs/indels in known cancer genes in the pathogenesis of these aggressive malignancies, but much has remained elusive about the breadth of other germline risk factors. By harmonizing 10,590 germline genomes, applying GATK-SV for comprehensive genome-wide SV discovery, implementing rigorous quality control, and leveraging an ancestry-aware case-control association framework, we show that rare germline SVs across the coding and noncoding genome are important risk factors for neuroblastoma, Ewing sarcoma, and osteosarcoma.

We found that large, unbalanced germline SVs were risk factors for all three pediatric solid tumor histologies. This signal was stratified by sex, with male cases having a four-fold higher incidence of these SVs than male controls while female cases exhibited rates comparable to female controls. Previous epidemiological studies have reported a slight male predominance across pediatric cancers, including the three histologies in our study (male-to-female incidence relative rates: 1.1-1.7),^60, 61^ but the reasons for this sex bias have remained opaque. The sex-specific pattern of large, unbalanced SVs we discovered may imply that there are specific categories of autosomal germline variants that differentially increase cancer risk in males and females, although additional studies are needed to unambiguously confirm this trend. In striking contrast to developmental disorders, where large CNVs at dozens of established loci are associated with disease risk,^35, 36^ the large unbalanced SVs observed in pediatric cancer cases did not coalesce at specific loci, and 91% did not overlap any genes with known roles in cancer. Instead, these SVs were seemingly scattered throughout the genome at random, hinting at a possible role in genomic instability during tumor initiation that echoes the previously reported associations of pathogenic germline rare variants in DNA damage repair genes in these same histologies.^47^ Similarly, the lack of any association with equally large but balanced SVs (e.g., inversions) was curious, as large inversions and translocations are strongly associated with other pediatric congenital anomalies.^34^ This discordance suggests that the pathogenic effects of large SVs in pediatric cancers are specific to dosage imbalances caused by exceptionally large germline CNVs.

Rare gene-disruptive germline SVs were also enriched in pediatric solid tumor cases relative to adult controls, and this effect was most pronounced for ultra-rare SVs. We implemented a comprehensive CWAS framework to identify specific subsets of SVs contributing to disease risk, which pinpointed LoF singleton SVs in mutationally constrained genes as an especially strong category of risk-carrying SVs in neuroblastoma. Further dissection of the CWAS results revealed nominally significant subsets of SVs with even greater effect sizes and higher biological specificity, such as SVs predicted to cause LoF of DNA damage repair genes in neuroblastoma and Ewing sarcoma. Mutationally constrained and DNA damage repair gene sets are therefore promising starting points for future discoveries of novel CPGs specific to pediatric solid tumors. We also found that risk-carrying SVs appeared to impact cellular functions with plausible connections to disease initiation, such as neurogenesis and synaptic transmission in neuroblastoma. By integrating our results with RNA data from normal tissues of adult donors and matched patient tumors, we further found that these risk-carrying SVs tended to affect genes that were highly expressed in the disease’s putative tissue-of-origin and even led to altered expression in the eventual tumors from SV-carrying patients.

We identified dozens of germline SVs impacting established cancer genes and pathways, such as *PHOX2B, BARD1, MYCN*, and RAS-MAPK genes in neuroblastoma or DNA damage repair/Fanconi Anemia genes in Ewing sarcoma, which demonstrates that germline SVs converge onto many of the same oncogenic genes as germline SNVs/indels and somatic alterations. However, the rate of rare gene-disruptive germline SVs across all known CPGs and COSMIC cancer genes was not significantly higher in cases relative to controls, bolstering the argument along with our CWAS results that there are likely CPGs specific to these pediatric malignancies that have not yet been discovered. Our gene-based rare SV association tests identified just one novel candidate CPG: LoF of *KL* in neuroblastoma, which may plausibly function as a tumor suppressor based on existing experimental evidence^53, 54^ but requires replication and further functional evaluation in these contexts. The robust identification of yet more undiscovered CPGs will likely require greater power afforded by the integration of SNVs, indels, and SVs in larger cohorts.

Lastly, we tackled the open question of how noncoding germline SVs may confer disease risk. By incorporating tissue-of-origin epigenetic annotations into our CWAS framework, we discovered that rare noncoding SVs overlapped TAD boundaries in adrenal gland tissue more frequently in neuroblastoma cases than controls. Deletions of TAD boundaries close to adrenally-expressed constrained genes had larger effect sizes than TAD boundary SVs generally, suggesting that dysregulation of TADs specifically affecting developmental genes expressed in the putative tissue-of-origin may be important in neuroblastoma pathogenesis. The lack of significant noncoding SV categories in Ewing sarcoma was intriguing, as comparable sample sizes implies that rare noncoding germline SVs play a larger role in neuroblastoma oncogenesis, although we cannot rule out effects specific to certain cell types or developmental timepoints that are not represented in existing public datasets.

Our study had notable limitations. Like most rare disease research, the scarcity of large patient datasets limited our power for gene- and locus-level association tests and prevented replication of our findings in an independent dataset. We also did not focus on common SVs in this study as their effect sizes are expected to be significantly smaller and require greater sample sizes than rare variant analyses. Lastly, our noncoding and functional analyses relied on existing datasets derived from healthy adult tissues or matched tumors of patients from this study, neither of which are perfect proxies for healthy fetal or pediatric tissues matched on developmental timing. Ongoing germline WGS efforts in other pediatric cancer populations will enable further generalization and replication of our findings, as well as the discovery of novel disease-specific risk factors.

In conclusion, this work represents a substantial step forward in explaining the missing heritability of pediatric solid tumors through the systematic characterization of germline SVs at nucleotide resolution. Our results lay the groundwork for the eventual incorporation of germline SV analyses into routine diagnostic screening and clinical practice. Future work focused on integrated genome-wide evaluation of SVs with SNVs/indels in the same patients will enable a more complete understanding of germline factors contributing to the development of pediatric cancer, and in turn a better understanding of the central oncogenic programs giving rise to these malignancies.

## Acknowledgements

We gratefully acknowledge the participants and studies who provided biological samples to enable this research. We thank Dr. Erica Pimenta for her thoughtful and critical review of this manuscript. We thank Drs. Joshua Schiffman, Schuyler O’Brien, Erin L. Young, Lucy Hayes, Gareth Mitchell, Trent Fowler, Jo Anson, and the team at the Huntsman Cancer Institute for their foundational work in establishing the GMKF Ewing sarcoma dataset.

We thank the Children’s Oncology Group, QuadW Foundation, Dr. Mark D. Krailo, and Dr. Donald A. Barkauskas for the provision of banked normal tissue from a case of Ewing sarcoma to enable PCR evaluation. The content is solely the responsibility of the authors and does not necessarily represent the official views of the National Institutes of Health.

## Author Contributions

Conceptualization: RG, RLC, EMVA. Formal analysis: RLC, RG, JC, JKJ. Funding acquisition: RG, KAJ, EMVA, KH. Investigation: RLC, RG, JC. Methodology: RLC, MW, A-SJ, CW, EP-H, MT, HB. Resources: KAJ, BDC, JL. Supervision: EMVA. Validation: AG. Visualization: RLC, JC, RG. Writing – original draft: RG, RLC, JC. Writing – review & editing: EMVA, HB, AG, SHA, KAJ, BDC.

## Data & Code Availability

All raw genome sequencing data analyzed in this study is hosted by dbGaP (cohorts: MESA, BioMe, GMKF), St. Jude Cloud, the International Cancer Genome Consortium, or the International Genome Sample Resource (1000 Genomes Project; https://www.internationalgenome.org). Accession numbers for each cohort are provided in **Materials and Methods** and **Table S1**. The GATK-SV pipeline is publicly available from GitHub (https://github.com/broadinstitute/gatk-sv). All other software and code used to generate the results in this study is publicly available from GitHub (https://github.com/vanallenlab/ped_germline_SV).

