## Supplementary Information for "Rare germline structural variants increase risk for pediatric solid tumors"

Gillani & Collins et al.

Version 1 | April 2024

#### Table of Contents

##### *Materials & Methods*


Ethics approval & consent to participate 3
Sample selection 3
Single-sample SV preprocessing 3
Sample quality control, batching, sex inference, and read-depth CNV discovery 3
Joint SV filtering, clustering, and genotyping 4
Post hoc SV callset refinement 6
Functional consequence annotation 8
Ancestry & relatedness inference 9
Ancestry rebalancing & sample pruning 9
SV frequency annotation & definitions for rare SVs 10
Final SV callset benchmarking & quality assessments 10
Case-control association testing 11
Genome-wide sliding window association tests of large, rare CNVs 11
Coding category-wide SV association study framework 12
Gene-disruptive events in germline cancer predisposition genes and COSMIC cancer genes 13
Gene-based rare SV association testing 13
Approach to noncoding CWAS 13
CWAS gene set enrichment 14
Expression dysregulation association with germline SVs 14


##### *Supplementary Text*


S1. SV dataset benchmarking. 16


##### *Supplementary Figures*


S1. WGS sample properties. 17
S2. Principal component analysis of common SVs. 18
S3. Ancestry-aware SV quality assessments. 19
S4. Supporting results for analyses of large, unbalanced SVs. 20
S5. Example read depth profiles for large, rare, unbalanced SVs in pediatric cancer cases and adult controls. 21
S6. Genome-wide rare CNV association scans. 23
S7. Supporting results for gene-centric analyses. 24
S8. Gene set enrichment and expression dysregulation associated with ultra-rare germline SVs in Ewing sarcoma and neuroblastoma. 25
S9. Rare coding SVs impacting established cancer genes. 26

S10. Lack of enrichment of rare germline SVs at sites of recurrent somatic SVs in Ewing sarcoma and neuroblastoma. 27
S11. Gene-based coding SV burden tests. 28
S12. Noncoding CWAS schematic. 29
S13. Results of PCR in banked white blood cell DNA from a Ewing sarcoma case with an EWSR1-FLI1 translocation. 30

***Supplementary Tables*** 31

***Supplementary References*** 33

### Materials & Methods

#### Ethics approval & consent to participate

Written informed consent from individuals and institutional review board approval, allowing comprehensive genetic analysis of germline samples, were obtained by the original studies that enrolled individuals. The secondary genomic analyses performed for this study were approved under Dana-Farber Cancer Institute institutional review board protocols 21-143 and 20-691. This study conforms to the Declaration of Helsinki.

#### Sample selection

Germline WGS data from neuroblastoma (NBL), Ewing sarcoma (EWS), and osteosarcoma (OS) cases were aggregated across the Gabriella Miller Kids First (GMKF; dbGaP study "Genomic Sequencing of Ewing Sarcoma" phs000804.v1.p1 with individuals enrolled through Project GENESIS at the Huntsman Cancer Institute and Children's Oncology Group protocol AEPI10N5; dbGaP study "Discovering the Genetic Basis of Human Neuroblastoma: A Kids First Project" phs001436.v1.p1), St. Jude Cloud (Pediatric Cancer Genome Project, St. Jude Lifetime, Genomes for Kids, and Childhood Cancer Survivor Study), and International Cancer Genome Consortium (ICGC; "Bone Cancer – UK", BOCA-UK) studies. Germline WGS for a total of 2,277 cases (996 NBL, 887 EWS, 394 OS) were aggregated prior to quality control, outlier exclusion, and relatedness inference (*1-5*). Proband cases from the GMKF study were augmented with matched germline WGS from parents where complete parent-child trios were available (405 trios in NBL, 283 trios in EWS). 5,320 adult control samples were selected from the TOPMed BioMe (2,842; dbGaP study "NHLBI TOPMed - NHGRI CCDG: The BioMe Biobank at Mount Sinai" phs001644.v3.p2) and MESA studies (2,478; dbGaP study "NHLBI TOPMed: MESA and MESA Family AA-CAC" phs001416.v3.p1) (*6*). Controls were selected based on reported ancestry to approximately match the proportions of genetic ancestry among NBL, EWS, and OS cases to enable maximally powered ancestry-matched association analyses. Finally, 539 parent-proband WGS trios (1,617 individuals) from the 1000 Genomes Project (1000G) were included for batched analysis with the selected cases and controls to enable downstream quality control, SV benchmarking, and robust ancestry inference (*7*). Across these cohorts, WGS was performed on DNA extracted from predominantly three main sources, including peripheral blood, saliva, or—in the case of 1000G samples only—lymphoblastoid cell lines (**Fig. S1**).

#### Single-sample SV preprocessing

All WGS samples were processed according to the recommended default configuration of GATK-SV v0.24.3-beta, which has been described in detail in previous publications and is available from GitHub (see Data and materials availability) (*8*). The initial single-sample step of GATK-SV executes three complementary SV discovery algorithms in parallel for each WGS sample: Manta v1.5.0 (*9*), MELT v2.0.5 (*10*), and Wham v1.7.0-311-g4e8c (*11*). Additionally, raw SV evidence metrics were collected genome-wide for each WGS sample across four evidence classes, including anomalous paired-end (PE) reads, split reads (SR), read depth (RD), and B-allele frequencies (BAF) of SNVs and indels.

#### Sample quality control, batching, sex inference, and read-depth CNV discovery

The first half of the GATK-SV pipeline operates on batches of 100-1,000 samples. We therefore undertook a joint batching and sample quality control process to simultaneously exclude low-quality WGS samples while dividing our cohort into homogenous batches of samples for subsequent SV discovery & genotyping. This procedure roughly follows the protocol used for the most recent release of the Genome

Aggregation Database (gnomAD) (*8*). We began by excluding extreme outlier WGS samples whose technical qualities deviated markedly from all other samples. We defined these "global" outlier samples as those meeting any of the following criteria:

- Dosage bias (∂; i.e., coverage nonuniformity) less than the first quartile (Q1) minus eight times the median absolute deviation (MAD) across all samples;
- ∂ greater than the third quartile (Q3) plus eight MAD across all samples;
- Median WGS coverage <15 or >60;
- WGS insert size < Q1 – 8*MAD;
- WGS insert size > Q3 + 8*MAD;
- Highly variable autosomal ploidy estimates, defined as the median absolute copy number deviation from diploid exceeding >0.1 across all 22 autosomes;
- Ambiguous sex chromosome ploidy;
- Inferred XX or XY karyotype that disagreed with a self-reported binary sex (i.e., inferred XX but reported "male" or inferred XY but reported "female").

In total, 364 samples (3.4% of total cohort) failed one or more of the criteria above and were excluded outright from the study.

Following global outlier exclusion, we inferred sex chromosome ploidy for all samples using GATK-SV's ploidy inference functionality (e.g., **Fig. 2B**; **Fig. S4A-B**) before dividing the remaining 10,226 samples into 20 batches of 509-513 samples per batch. We assigned samples to batches roughly matched on ∂ and genome-wide median coverage while ensuring that all 20 batches also had the same proportions of male cases, female cases, male controls, and female controls. Once all samples were assigned to a batch, we performed a second layer of outlier exclusion within each batch. We excluded samples from each batch if they violated any of the following criteria, all of which were computed while restricting to samples in a given batch:

- ∂ < Q1 - 7*MAD;
- ∂ > Q3 + 7*MAD;
- WGS insert size < Q1 - 7*MAD;
- WGS insert size > Q3 + 7*MAD.

A total of 77 additional samples failed these criteria; these samples were removed from their batch and excluded outright from the remainder of the study.

This joint batching and outlier exclusion procedure retained a total of 10,149 samples across 20 batches of 491-513 samples per batch. Following batch assignment, we conducted RD-based CNV discovery for all samples per batch using GATK-gCNV v4.2.6.1 (*12*) and cn.MOPS v1.40.0 (*13*), both of which were executed using GATK-SV recommended default parameters. The CNV segments predicted by GATK-gCNV and GATK-SV were merged per sample using GATK-SV default parameters to generate a single set of RD-based CNV predictions for each sample.

#### Joint SV filtering, clustering, and genotyping

We used GATK-SV to perform joint SV filtering and genotyping across all 10,149 samples passing initial quality thresholds. This pipeline has been extensively detailed in prior studies (*8*) and all code and parameters are available from GitHub. Accordingly, only an overview is outlined below; please refer to the GATK-SV documentation and source code for exact technical specifics. The GATK-SV pipeline involves a series of operations for each batch of samples prior to integrating the results per batch across all batches in a cohort. We performed the following operations per batch in the order listed below:

1. SV calls from each algorithm were clustered across all samples in a batch using default clustering GATK-SV parameters.
2. Raw WGS evidence (i.e., PE, SR, RD, BAF) was collected for all samples for each candidate SV from step 1.
3. A series of random forest classifiers were trained on the output of step 2 to predict true positive SVs and exclude technical false positives. After training, these classifiers were applied to all SVs from step 1 and only predicted true positive SVs were retained for subsequent steps.
4. Outlier samples were excluded from each batch based on SV counts after SV filtering from step 3. Outliers were defined as samples with SV counts either < Q1–(10*MAD) or > Q3+(10*MAD). Outliers were assessed separately for every combination of algorithm (e.g., Manta) and SV type (e.g., deletion), and any sample flagged as an outlier for any of these strata was excluded from the batch and from the remainder of the study. In total, 216 (2.1%) of all samples were excluded at this stage; the remaining 9,933 samples were retained for subsequent processing and analysis.
5. SV calls from the three PE/SR-based algorithms—Manta, MELT, and Wham—were clustered within each batch to produce a single set of PE/SR-based SV calls for each batch.

After completing the above operations per batch, we next unified the SV predictions from each batch across all samples in the cohort to produce a uniformly genotyped SV callset for the entire cohort. This required several sequential operations performed cohort-wide, as follows:

6. A nonredundant list of all candidate SVs across all batches was constructed separately for PE/SR- and RD-based SV calls. No clustering was performed in this step.
7. Each sample was assigned a genotype and genotype quality (GQ) score for every candidate SV present in the output from step 6. These genotypes were assigned using GATK-SV's default SV genotyping module, which reuses the SV evidence thresholds learned by the random forest classifiers from step 3 to predict biallelic genotypes based on the strength of evidence supporting each SV.
8. After genotyping, SVs were clustered across all batches using default GATK-SV parameters. This algorithm first clusters PE/SR- and RD-based SVs separately across all batches before subsequently clustering the cohort-wide PE/SR- and RD-based SVs into a single unified file of nonredundant, clustered SVs with genotypes for every sample.
9. Complex SVs were resolved using the standard GATK-SV approach, which involves first identifying clusters of proximal SV breakpoints in the same sample(s) that may represent complex SVs before comparing the coordinates of these breakpoints to a reference taxonomy of 16 possible germline complex SV allele configurations using svtk resolve (*8, 14, 15*). Candidate complex SVs involving predicted segments of deletion or duplication were further interrogated by re-genotyping the predicted CNV interval(s) based on RD evidence for all predicted carrier samples; complex SVs that were not orthogonally confirmed by RD re-genotyping were excluded as "unresolved" variants.
10. The final module of the core GATK-SV pipeline, "CleanVCF", applied a series of minor adjustments to handle myriad edge cases, such as adjusting genotypes for male and female samples on allosomes, detecting and re-genotyping multiallic CNVs, and assessing predicted copy numbers for samples with two independent overlapping CNVs (i.e., predicted compound heterozygotes).

The final output of the core GATK-SV pipeline was a single VCF comprising all SVs detected in the cohort with predicted genotypes for all 9,933 samples.

#### *Post hoc* SV callset refinement

Following joint SV discovery and genotyping across all samples, we performed a series of *post hoc* cleanup steps to further improve the specificity of our SV calls and genotypes. We largely followed examples set by gold-standard GATK-SV callsets generated by gnomAD and the NIH All of Us Project (https://support.researchallofus.org/hc/en-us/articles/14941865780500-Benchmarking-and-quality-analyses-on-the-All-of-Us-short-read-structural-variant-calls) (*8, 16*); we deviated slightly in some respects to emphasize specificity at the expense of sensitivity, reasoning that spurious false positives were less desirable than false negatives given our focus on rare variant associations in rare diseases. In total, these cleanup steps involved a mixture of established and new methods applied in the following order:

1. We first pruned low-quality genotypes using a pre-trained model, FilterGenotypes, that was developed as an extension for GATK-SV in the context of the NIH All Of Us WGS dataset (*16*). Detailed documentation for the Genotypes model is available from the All of Us genomic data quality report available from the All of Us online (https://support.researchallofus.org/hc/en-us/articles/4617899955092-All-of-Us-Genomic-Quality-Report; see pages 34-37). In brief, FilterGenotypes computes a scaled likelihood (SL) for each genotype, which is proportional to the log-odds that the genotype is a true positive versus a false positive, then optimizes minimum SL thresholds below which genotypes are deemed low-confidence. Low-confidence genotypes are erased and reassigned to no-calls (i.e., "./." in VCF). The FilterGenotypes model was trained on GATK-SV data for 606 All of Us participants to optimize concordance with long-read WGS and SNP microarrays performed on the same set of 606 participants. We applied the pre-trained "high sensitivity" FilterGenotypes model from All of Us directly to the SV callset we generated in the present study. The SL cutoffs from the "high sensitivity" model corresponded to an approximate 15% false discovery rate (FDR) in All of Us according to publicly available All of Us benchmarking information (see URL above).
2. We further excluded residual low-quality genotypes not captured in step 1 by applying a second method, minGQ, which was developed by gnomAD to optimize genotype quality thresholds based on empirical rates of SV transmission between parents and children in complete trio pedigrees (*8*). The minGQ method learns a decision tree of minimum GQ thresholds to retain genotypes given an SV's size, frequency, type, and other technical features. We trained a minGQ filtering model to optimize cutoffs for a false discovery rate of 5% for the pediatric cancer parent-child trios from GMKF that were present in our dataset at this stage.
3. All multiallelic CNVs (mCNVs) smaller than 5kb were excluded from our final callset. We have previously observed that mCNV predictions from GATK-SV below this size threshold are often unreliable (*8*).
4. We implemented a procedure to identify splice junctions from genic retrocopy insertion polymorphisms (GRIPs), which can appear in WGS as deletion SVs precisely spanning exon-exon junctions despite not corresponding to real deletions of endogenous DNA (*17*). We systematically detected these spurious deletion calls by intersecting all deletions lacking RD support with introns defined in the MANE Select v1.2 human gene reference catalog (*18*). Any such deletion with ≥95% reciprocal overlap with an annotated intron was tagged as a likely GRIP splice junction and not included for any downstream analyses.
5. Common SVs (AF>5%) that were predicted to be >500kb in size were marked as "unresolved" variants; these edge-case SV predictions usually correspond to small polymorphic insertions that were incorrectly resolved as large SVs by GATK-SV.
6. Common reciprocal translocations (AF>1%) were marked as "unresolved" for the same reason as step 5.

7. Any SV that exhibited >20% coverage by loci that had at least one alternate haplotype in the hg38 reference assembly were excluded to forestall possible issues related to variant callers handling multiply mapped reads corresponding to alternative haplotypes.
8. Genotypes for each sample were masked if they had GQ of zero prior to step 1 (FilterGenotypes) and corresponded to uncommon (AF<5%), biallelic SVs called exclusively by Manta or Wham.
9. SVs were excluded if they exhibited a mean SL<0 and max SL<75 across all non-reference samples, were called exclusively by Wham or Manta, were smaller than 1kb in size, did not exhibit BAF evidence, were uncommon (AF<5%), had a no-call genotype rate (NCR; i.e., rate of pruned or missing genotypes) >0.1%, and did not exhibit overdispersed PE/SR genotyping evidence.
10. We applied a final layer of systematic outlier sample exclusion. For each sample, we collected counts of (i) all SVs, (ii) all SVs per SV type, (iii) all rare SVs, (iv) rare SVs per SV type, (v) deletions with AF<5% between 400bp and 1kb in size, (vi) CNVs overlapping transcription start sites, and (vii) CNVs overlapping protein-coding exons. For each strata above, we defined outlier samples on a population-specific basis as samples with SV counts less than Q1 – 5*MAD or greater than Q3 + 5*MAD within each population based on preexisting ancestry estimates (*19, 20*) and/or self-reported race and ethnicity information if necessary. Samples with unknown or nonspecific ancestry or self-reported race/ethnicity were grouped with European samples for the purposes of this analysis. All SV count strata were treated equally when determining outliers except for strata where all samples had <10 SVs or the population-wide MAD of SV counts was zero; such especially sparse strata were not considered for this outlier definition. Any sample labeled as an outlier in any of the strata or populations above was excluded outright from the study and not included in any subsequent analyses. In total, this process excluded 557 samples, leaving 9,377 non-outlier samples in our overall cohort for downstream steps.
11. Any SV with NCR≥4% was excluded. This threshold was decreased to NCR≥0.4% for deletions between 400bp and 1kb in size with AF<5%. All samples were included for NCR calculations on autosomes, whereas we only considered female or males samples when defining NCRs on chromosome X or Y, respectively.
12. We excluded 15 CNVs ≥1Mb with AF<2% after manual review of RD evidence for each CNV as follows. We visualized RD profiles for all 121 biallelic CNVs ≥1Mb with AF<2% using standard visualization approaches implemented in GATK-SV (*15, 21*). We found strong visual support in RD profiles for 106 CNVs that were consistent with a germline gain or loss of a full copy (e.g., **Fig. S5**). We noted an additional two CNVs that had clear visual RD support but at a sub-integer copy number, potentially indicative of post-zygotic mosaic mutation origins and/or sample contamination. Finally, we found no clear RD support for 13 CNVs, concluding that these were technical false positives that had been incorrectly resolved as very large CNVs by GATK-SV. To remain conservative, we retained only the 106 CNVs that exhibited strong integer copy states consistent with germline mutational origins based on RD evidence; the other 15/121 CNVs were excluded from the remainder of the study.
13. We excluded four complex SVs based on a manual review of RD profiles similar to step 11. We reviewed all 19 complex SVs that were ≥1Mb in size, AF<2%, and had at least one predicted CNV segment ≥5kb. We found strong integer copy-number support for 15/19 and found little-to-no convincing evidence for 4/19. We did not observe any candidate mosaic SVs in this review.
14. SVs were excluded if they exhibited significant AF bias between pairs of case or control cohorts (e.g., St. Jude vs. GMKF or MESA vs. BioMe). We performed a Fisher's exact test of allele counts for pairs of cohorts for each SV with AF<5% in our overall dataset and excluded any SVs with P<0.05. This test was further controlled for genetic ancestry and population stratification on a variant-by-variant basis by restricting each test to the continental ancestry group with the largest number of variant

carriers as has been suggested previously (*8*). We did not at any point compare cases and controls in this analysis to avoid confounding technical batch effects with potentially disease-associated SVs.

15. We masked all genotypes for CNVs exclusively with RD evidence ("RD-only CNVs") in samples with unusually high RD-only CNV counts. We first collected counts of RD-only biallelic deletions and duplications in all samples and defined a sample as an RD-specific outlier if their count of RD-only deletions or duplications was >Q3 + 5*MAD within their respective continental ancestry group. These samples were retained in our overall dataset as their counts of non-RD-only SVs were well-behaved, but we masked the genotypes of RD-only samples to prevent these samples from disproportionately contributing noise to analyses including RD-only CNVs.

16. Lastly, three samples with canonical oncogenic translocations (one control with *BCR-ABL1*, one Ewing sarcoma case with *NUP98-PRRX1*, and one Ewing sarcoma case with *EWSR1-FLI1*) were assumed to be either tumor-normal sample swaps or significant tumor-in-normal contamination, and therefore excluded outright. To confirm that the *EWSR1-FLI1* translocation detected in the Ewing sarcoma case was not a germline event, we acquired a banked germline sample of white blood cells from the Children's Oncology Group and performed PCR to test for the predicted translocation breakpoints. Endpoint PCR was performed using Q5 High Fidelity DNA polymerase 2X Master Mix (New England Biolabs) in a 20 µl reaction volume consisting of 20 ng of DNA and 200 µM of each forward and reverse primer (IDT). The following primers were used:

17.

| Locus | Forward primer (5' to 3') | Reverse primer( 5' to 3') | Amplicon |
|---|---|---|---|
| *FLI1* (1) | GGGGCGGTGGTAATGGAG | TGAAACCACCACAAATGATGCT | 200bp |
| *FLI1* (2) | CATGCTTTGTCCACGCTTATCA | AGAAGATGTCTGAAGCCCGT | 117bp |
| *EWSR1* (1) | ATGGGCTCACTTCCTACTGGA | CCTTCTGGATTATGTTAACCACCA | 220bp |
| *EWSR1* (2) | CCAATAGCATTTTGCAGAGTAATGT | ACCTTCATCAAAGTACAGACACCA | 217bp |

The PCR reactions were performed in a C1000 Touch Thermal Cycler (Bio-Rad): initial denaturation at 98$^0$C for 30 seconds, followed by 30 cycles of 98$^0$C for 5 seconds, 64.3$^0$C for 30 seconds, 72$^0$C for 30 seconds, and a final extension at 72$^0$C for 2 minutes. PCR endpoint products were loaded in a 2% agarose gel with 1X TBE Buffer (ThermoScientific) at 75V for 1 hour. GelRed Nucleic Acid Stain (Biotium) was used to visualize the PCR product on Biorad Gel Dox XR+. Wild type *EWSR1* and *FLI1* sequences were amplified, but not the *EWSR1-FLI1* translocation sequence spanning the reported breakpoint (**Fig. S13**).

After applying all the 16 criteria above, we retained a total of 284,395 high-quality SVs carried by at least one of 9,374 high-quality samples. All subsequent analyses were restricted to this dataset. The code required to apply each of the steps above is freely provided on GitHub (see Data and materials availability).

#### Functional consequence annotation

We next annotated all predicted genic effects for each SV using GATK-SV's inbuilt annotation toolkit, "SVAnnotate". We used the MANE Select v1.2 gene reference dataset for all genic predictions (*18*). The approach used by GATK-SV to annotate genic consequences has been described previously (*8*). This method assigns one of 11 possible predicted consequences to each gene overlapped by each SV. For genes where more than one consequence could be assigned, more "severe" consequences are prioritized in a conceptually similar manner to other annotation frameworks for SNVs/indels (*22*) in the following order: loss-of-function (LoF), copy-gain (CG), intragenic exonic duplication (IED), partial exon duplication (PED), TSS duplication, partial gene duplication, spanning inversion, multiallelic SV exon overlap, UTR overlap, intronic, or promoter overlap. For the purposes of this study, we defined an SV as "gene-disruptive" if it was assigned a LoF, CG, IED, or PED consequence for any protein-coding gene.

Noncoding functional annotations were derived based on prior assays in the putative tissue-of-origin where available: adrenal gland in NBL and skeletal muscle in EWS. ATAC peaks, consensus enhancer tracks, TAD boundary definitions from HiC data, and H3K27Ac peaks were derived from ENCODE (*23*). Enhancer, genic enhancer, bivalent enhancer, and flanking active transcription start site annotations were used from the Roadmap Epigenomics Project 15-state ChromHMM model (*24*). Additional noncoding annotations included Activity-by-Contact model enhancers (Engreitz Lab) (*25*), ultraconserved noncoding elements (UCNE base) (*26*), recombination hotspots (deCODE from the UCSC Genome Browser) (*27*), and fragile sites (PCAWG consortium) (*28, 29*). Sources of noncoding annotations are detailed in **Table S8**. We annotated overlap between all noncoding annotation tracks and every SV in our dataset using GATK-SV SVAnnotate, which considers two possible relationships between SVs and noncoding elements: "breakpoint", wherein an SV breakpoint falls directly within the noncoding element, and "span", wherein the SV full spans the entirety of the noncoding element.

#### Ancestry & relatedness inference

We inferred genetic ancestry for all samples and relatedness between all pairs of samples using Hail v0.2.119. We first restricted to well-genotyped polymorphic autosomal SVs, which we defined here as autosomal SVs that had QUAL>10, NCR≤0.1%, 0.1%≤AF≤99.9%, and Hardy-Weinberg equilibrium (HWE) chi-square $P<10^{-5}$. We then computed the top 20 HWE-normalized principal components (PCs; **Fig. S2**) before inferring kinship coefficients and identity-by-descent proportions for all pairs of samples using Hail's implementation of PCRelate (*30*).

We assigned samples to one of five continental ancestry groups for the purposes of variant frequency annotation. To achieve this, we trained a linear SVM on the top 10 genetic PCs generated by Hail to cluster samples into the following five populations: African/African-American (AFR), admixed American/Latino (AMR), east Asian (EAS), European (EUR), and south Asian (SAS). We trained the SVM classifier on ground-truth population assignments available for samples from the 1000 Genomes project samples and supplemented with samples from the TOPMed MESA cohort that had previously undergone WGS SNV-based ancestry assignment as part of a prior study (*7, 20*). We validated the performance of our ancestry classifier by comparing its predicted labels to ancestry labels previously inferred from SNV/indel analyses for a subset of these samples not included in the training set (*19*). This ancestry classifier achieved accuracies of 98.5% and 97.1% on the training and validation sets, respectively, with classification errors primarily corresponding to admixed individuals that did not clearly cluster with any one ancestry group in PC1-10. Following training and validation, we applied this classifier to assign predicted genetic ancestry memberships to all samples in our cohort.

#### Ancestry rebalancing & sample pruning

Prior to undertaking SV-disease association analyses, we wanted to isolate a strictly unrelated set of ancestry-matched cases and controls for all association tests. We began by excluding all parents from known parent-child trios and all samples from the 1000 Genomes Project. We further pruned all predicted pairs of cryptic (i.e., unreported) first- or second-degree relatives from our dataset by (i) identifying all pairs of individuals with kinship coefficient ($\Phi$) ≥0.1, (ii) tallying the number of times each sample appeared in pairs of apparently related samples from step i, (iii) excluding the sample with the largest count of apparent relatives from step ii, and (iv) repeating steps ii-iii until no more predicted pairs of samples with Φ≥0.1 remained in our cohort. In the case of ties between two or more samples in step ii, we prioritized retaining affected cases and therefore preferentially removed controls where possible.

Next, within each disease (NBL, EWS, OS, or all histologies combined), we rebalanced the proportions of inferred genetic ancestries between cases and controls by iteratively downsampling controls from the ancestry groups with the largest absolute fold-difference in sample size between cases and controls until a Chi-square test of sample counts per ancestry between cases and controls converged to P<0.01. This process resulted in a set of 6,728 unrelated cases and controls with approximately equal representation from four major continental genetic ancestries, which we used for all subsequent disease association analyses in this study. Note that slight differences in ancestry composition between the three diseases required very slight deviations in sets of controls used for association testing per ancestry (control counts: NBL=4,830; EWS=4,574; OS=4,805; pan-cancer=4,717), although any sample used as a control for any disease was retained in our final dataset.

#### SV frequency annotation & definitions for rare SVs

As the final step prior to downstream association analyses, we used GATK-SV's annotation module to log the AFs of each SV in each of the five continental populations (AFR, AMR, EAS, EUR, SAS) based on the SV genotypes in our cohort, as well as a pooled estimate across all populations. All samples were included for autosomes; samples with sex chromosome abnormalities were excluded when calculating frequencies of SVs on allosomes. We further defined the cross-population maximum ("PopMax") AF as the maximum AF observed in any of the four populations with at least >100 samples in our final dataset (AFR, AMR, EAS, EUR). Additionally, we annotated the frequencies of the SVs detected in our cohort with the AFs of their corresponding SVs in gnomAD v3.0, if any (*8*). We recorded the SV AF from gnomAD for all major populations and further computed a gnomAD-specific PopMax AF across all non-bottlenecked populations with at least ≥100 samples included in gnomAD v3.0; these populations included AFR, AMR, EAS, non-Finnish EUR, and SAS.

Throughout the downstream analyses in this study, we frequently referred to two different sets of SVs defined by their frequency: "rare" and "singleton". The exact criteria for each of these SV sets is defined below:

- Rare: biallelic SVs with an in-cohort PopMax AF<1% after excluding first- and second-degree relatives, as well as gnomAD v3.0 PopMax AF<1% (*8*).
- Singleton: biallelic SVs with exactly one alternative allele observed across all unrelated samples in our dataset (i.e., AC=1 in the VCF generated in this study). We further required PopMax AF<1% in gnomAD v3.0 as for rare SVs, reasoning that many singletons in our cohort would likely represent ultra-rare polymorphisms in the general population and therefore we were intentionally more permissive with gnomAD frequency criteria for singletons in our cohort.

#### Final SV callset benchmarking & quality assessments

We performed four analyses to assess the technical quality of our final SV dataset (**Note S1; Fig. S3**). The specifics of each analysis are provided below:

1. Frequency comparisons to gnomAD: we assessed the concordance between the AFs of SVs detected in our study and their corresponding AFs in gnomAD v3.0 (*8*) using the same gnomAD AF annotations described above. We performed Pearson correlation tests for SV AFs within each of five continental ancestries that were present in gnomAD and in our study. The one exception was Europeans: given that gnomAD annotates Finnish and non-Finnish Europeans (NFE) separately, we compared gnomAD NFE AFs to the EUR AFs in our study.

2. Hardy-Weinberg Equilibrium: we tested whether the genotype distribution for each autosomal, biallelic SV was consistent with expectations under HWE. For each SV, we performed a Chi-square test of genotype counts for each of the five major continental ancestry groups considered in our study using the R HardyWeinberg package v1.7.5 (*31*). We considered an SV to be significantly deviant from HWE if the Chi-square P value surpassed a Bonferroni significance threshold after accounting for all SVs tested within a given continental ancestry group.
3. Mendelian transmission: we evaluated the rate of SV inheritance in 859 parent-child trios where all three family members had passed all filters and were present in our final SV callset. We first documented the biallelic SVs with non-reference genotypes in each child's genome and annotated each SV with the genotypes for both parents, only retaining sites with informative (i.e., non-missing) genotypes in all three family members. Next, for each family, we computed the fraction of SVs present in the child genome that were not found in either parent; this fraction reflects a combination of false positive genotypes in the child and false negative genotypes in one or both parents. Lastly, we computed the median of these fractions across all 859 trios as the overall assessment for the cohort.
4. Comparisons to long-read WGS: a subset of 24 samples from the 1000 Genomes project included in our study also had germline SV calls based on long-read genome assemblies from the Human Genome Structural Variation Consortium (HGSVC) (*32*). We leveraged these data to assess the precision of the short-read-based SV calls in our study. We applied GATK-SV module 09 to identify matching SVs in our short-read dataset and the HGSVC long-read dataset. This approach has been described elsewhere (*7, 8*), but in brief it searches for overlapping SVs present in the same samples and applies additional criteria based on SV size and type; generally, smaller SVs (<5kb) are considered a match based on single-linkage clustering of their breakpoints (±250bp), whereas larger SVs (≥5kb) are not required to have breakpoint clustering but instead must exhibit 50% reciprocal overlap. Given that short-read SV calls are less accurate in the most highly repetitive genomic regions (*33*), we restricted this analysis to SVs that did not have more than 10% overlap with annotated segmental duplications or simple repeats in the hg38 primary assembly. After excluding SVs in repetitive loci, we computed the fraction of short-read SVs per genome that were supported by long-read SVs in that same sample and reported the median across all samples as the overall summary metric for the cohort.

#### Case-control association testing

Unless otherwise specified, all case-control association tests reported in this study correspond to a logistic general linear model (GLM) with affected status (case|control) as the outcome variable, count of SVs as the primary predictor variable, and covariates for sample sex, cohort, and genetic PCs 1-3. All cases and their corresponding ancestry-matched controls were included except for samples that were missing >5% of SV genotypes for that test (e.g., samples missing genotypes for >2 SVs out of a set of 40 SVs would be excluded). For tests with extremely sparse SV data causing the GLM to fail to converge or produce coefficients with unbelievably large standard errors (log-odds standard error >10), we reran the association test using Firth's bias-reduced penalized logistic regression as has been implemented in many gold-standard tools for conventional genetic association studies of SNVs and indels (*34, 35*).

#### Genome-wide sliding window association tests of large, rare CNVs

We scanned all autosomes for loci that exhibited excesses of large (≥100kb), rare germline SVs in pediatric cancer cases vs. adult controls. We conducted these tests using a sliding window approach as has been recently proposed in similar studies of large, rare CNVs in developmental disorders (*36*). We first segmented all 22 autosomes into 1Mb windows sliding across each contig in 250kb steps before excluding any windows with ≥30% coverage by segmental duplications, simple repeats, gaps in the reference assembly, loci with alternative haplotype contigs, and loci with reference patches. For each window, we next identified all rare, large SVs involving at least ≥100kb of deletion or duplication that overlapped the window. We performed one case-control association test per window for each combination phenotype (NBL, EWS, OS, or pan-cancer) and CNV type (deletions only, duplications only, or all CNVs), resulting in a total of 12 tests per window (**Fig. S6**). We assessed significance after correcting for the total number of non-overlapping 1Mb windows tested (N=2,623.5 independent windows; $P<1.9\times10^{-5}$).

#### Coding category-wide SV association study framework

To systematically evaluate the differential burden of groups of SVs impacting genes in cases relative to controls, we carried out a "category-wide association study" (CWAS) of gene-disruptive SVs in EWS and NBL (*15, 37*). We enumerated 4,800 possible coding SV categories per disease based on six combinatorial layers of filters: SV AF, predicted coding consequence, SV type, genic mutational constraint (*38*), gene set membership (*39*), and tissue-of-origin gene expression (*40*) (**Tables S3-S5**). For each filter layer, we also included a catch-all "any" filter value that included all SVs. Loss-of-function constrained genes were defined as genes in the top sextile of the LOEUF metric as reported in gnomAD, whereas missense constrained genes were defined as the top sextile of missense observed/expected (*38*). All protein-coding genes were defined according to MANE Select v1.2 (*18*), and functional gene sets were derived from the Reactome Pathway Database. Tissue-of-origin expressed genes were defined as the subset of genes with a minimum expression of TPM>5 in adrenal gland and skeletal muscle from the Genotype-Tissue Expression project (GTEx) for NBL and EWS, respectively (*40*). Deletion, duplication, inversion, complex, and insertion SVs with predicted gene-disruptive consequences (copy gain, loss-of-function, intragenic exon duplication, and partial exon duplication) located on autosomes were assigned to coding variant categories based on SV and gene properties. To focus on categories for which we had sufficiently dense SV counts, we restricted all analyses to categories meeting a minimum threshold of >10 SVs across cases and controls; this yielded 714 and 679 "testable" categories in NBL and EWS, respectively. Case-control burden tests for each category were performed using the logistic GLM as described above.

Many categories included overlapping sets of SVs and were thus highly correlated. To estimate the effective number of independent tests performed in each histology, sample IDs were randomly permuted relative to SV counts 1,000 times for each category, and burden testing was repeated for each permutation to generate simulated Z scores. A Pearson correlation matrix between categories based on permuted Z scores was generated, and this correlation matrix was decomposed into eigenvalues and eigenvectors. We estimated the effective number of independent tests conducted for each disease to be the number of eigenvalues that captured 99% of variance in the inter-category correlation matrix. This estimated number of effective tests was used for the Bonferroni correction in each disease, resulting in a multiple hypothesis testing threshold of $\sim1.6\times10^{-4}$ for NBL and EWS.

To visualize the relationships between categories, we depicted them as a graph with nodes representing categories and edges representing filters (e.g., **Fig. 4A**, **Fig. 6E**). Categories were connected if the application of a filter to one category resulted in the other (e.g., a "deletion" filter to "all singletons" yields "all singleton deletions"). To depict the high degree of similarity between some categories, the y positions of the categories were then adjusted using a force-directed approach in which the force was proportional to their Jaccard similarity via SV overlap, such that similar categories were pulled together. To reduce the number of categories depicted, categories with Jaccard similarity >0.40 were collapsed, keeping the one with the lower P value and inheriting the other's edges.

#### Gene-disruptive events in germline cancer predisposition genes and COSMIC cancer genes

We undertook manual review of a focused subset of ultra-rare (AF<0.1%) gene-disruptive SVs in select germline cancer predisposition and COSMIC cancer genes (148 established cancer predisposition genes and 7 additional somatic drivers in NBL, EWS, and OS; total N=155 genes; **Table S3**). To maximize sensitivity, we began by identifying SVs intersecting the exons of the gene set above across all 1,992 cases in our intermediate SV callset prior to quality control and outlier exclusion. Candidate SVs were visualized and manually reviewed using a combination of the Integrative Genomics Viewer (*41*) and GATK-SV's native RD visualization functionality employed in the same manner as for the large CNV review during SV callset quality control (see above) (*8, 21*). SVs with a visually unambiguous accumulation of evidence upon manual review were deemed to be true positives for the purposes of this clinically focused analysis. Matched tumor copy number analysis was carried out using GATK v4.0.5.1 for the case identified as having a *de novo* germline *MYCN* duplication (*42*). Tumor-in-normal contamination was assessed using deTiN v1.8.5 (*43*). Separately, we quantified overall rates of rare and singleton gene-disruptive SVs in our final high-quality SV dataset (i.e., not only restricted to manually reviewed variants described above) that impacted a set of cancer predisposition genes (N=148) or COSMIC Cancer Census Tier 1 genes (N=566; **Table S3**) in NBL, EWS, and osteosarcoma cases compared to controls.

#### Gene-based rare SV association testing

We performed case-control rare SV association tests for 15,544 autosomal protein-coding genes annotated in MANE Select v1.2 whose gene bodies had <10% overlap with segmental duplications, simple repeats, reference patch loci, unalignable reference loci, and loci with alternative haplotype contigs in the hg38 assembly. For each disease, we separately evaluated rare gene-disruptive SVs (defined above), rare LoF SVs, and rare CG SVs using the logistic GLM approach as in all other analyses. Genes were considered significant if they reached a Bonferroni-adjusted threshold accounting for the number of genes tested ($P<3.2\times10^{-6}$).

#### Approach to noncoding CWAS

We extended our coding CWAS framework described above to focus exclusively on SVs with no predicted coding consequence with an additional functional layer capturing epigenetic and other noncoding genomic annotations (**Tables S8-10**). We enumerated 26,400 possible noncoding SV categories per disease based on 8 combinatorial layers of filters: SV AF, genic relationship (e.g., intronic, promoter, UTR, intergenic), SV type, functional element overlap, nature of intersection with the functional element, mutational constraint of the nearest gene, gene set membership of the nearest gene, and tissue-of-origin expression of the nearest gene. Deletion, duplication, inversion, complex, and insertion SVs with no known coding consequences and locations on autosomes were assigned to noncoding SV categories accordingly. Consistent with our previous coding CWAS, we again restricted to categories meeting a minimum threshold of >10 SVs across cases and controls; this yielded 7,909 and 6,244 "testable" categories in NBL and EWS, respectively. We once again evaluated the burden of autosomal SVs in cases relative to controls and identified noncoding categories meeting Bonferroni significance thresholds as described above, with a multiple hypothesis testing threshold of $\sim5.4\times10^{-5}$ for NBL and EWS. Finally, saddlepoint approximation was used to recalibrate the null distribution used for inference, from which we subsequently recalibrated P values and effect size point estimates (*44, 45*).

#### CWAS gene set enrichment

We used an approach similar to GO-term analysis to identify associations between CWAS categories and gene sets, modified to use SV counts. We first downloaded biological process gene sets from GO (*46*). Recognizing that we would have little power to detect enrichment for small gene sets and that large gene sets are usually nonspecific, we included gene sets containing between 30 and 1000 genes (N=2,722). The association between category and gene set based on SV counts was calculated using a Fisher's exact test with the following contingency table:

| | |
|---|---|
| Count of gene-set genes affected by category SVs | Count of non-gene-set genes affected by category SVs |
| Count of gene-set genes affected by non-category SVs | Count of non-gene-set genes affected by non-category SVs |

A continuity correction of one was added to all cells, and the "background" of non-category SVs was defined as all rare SVs that affected genes. For each analysis, this background was further filtered such that (i) the background SV class matched the category of interest (coding vs. noncoding), (ii) SVs affecting more than 10 genes were removed, (iii) SVs that did not have a coding/noncoding consequence detailed in the CWAS were removed, and (iv) noncoding SVs farther than 500 kb from their nearest gene for the noncoding consequence "INTERGENIC" were removed.

This Fisher's exact test was calculated for all gene sets, all Bonferroni-significant categories, and for cases and controls separately. We note that some categories–particularly those with functional filters like "affected gene expressed in adrenal gland"–can have substantial enrichment for certain gene sets even under a null of random SV locations because these categories are filtered to specific subsets of the genome. We therefore focused most of our discussion on "high-level categories" without functional filters, like "all singleton deletions". We also note that this contingency table ignores genomic covariates that influence which genes are affected by SVs (like gene size), so we relied on the control cohort, which will share the same technical confounders, for comparison to cases.

#### Expression dysregulation association with germline SVs

We used GTEx v8 to evaluate the normal tissue expression of genes affected by rare germline SVs in our cohort (*40*). We used the matrix of median TPMs by tissues to define expression of each gene in each tissue, focusing on adrenal gland and skeletal muscle for NBL and EWS.

To evaluate the effect of rare germline SVs on tumor expression, we leveraged RNA sequencing from NBL tumors available for a subset of the GMKF cohort (n=89/688) (dbGaP study "Discovering the Genetic Basis of Human Neuroblastoma: A Kids First Project" phs001436.v1.p1). We assessed the effect on expression of a group of SVs by category.

We converted sample TPMs to z-scores (for visualization in **Fig. 4E**), and then to sample ranks (0-88, normalized to 0-1). If a group of SVs does not affect expression of their associated genes, sample-normalized expression ranks will be uniformly distributed between 0 and 1. To account for positive or negative effects on expression (e.g., copy gain/deletions), we instead considered the absolute rank deviation from 0.5 (i.e., |0.5 - normalized rank|), in which case the null is uniform(0, 0.5), which has mean 0.25 and variance 1/48. We defined our test statistic as the mean absolute rank deviation over the samples with SVs in a group. Because the mean of n samples from any distribution is normal by the central limit theorem, we could directly calculate a P value for an observed mean absolute rank deviation $r$ with $n$ samples with SVs using the following formula:

$$\text{two-sided P value} = 2P(X < r \mid X \sim \text{norm}(0.25, 1/48n))$$

We assessed the Bonferroni-significant categories for NBL as well as larger groups like “all coding SVs”. To compare the effect of singleton SVs affecting genes expressed in adrenal tissue (n=25 affected genes) to any singleton coding SV (n=233 affected genes), we bootstrapped a P value by sampling 25 values from the “any coding SV” group ten million times and calculated how often the observed rank mean was less than that of the singleton SVs affecting adrenal genes group.

#### Supplementary Text

**Note S1 | SV dataset benchmarking.** We benchmarked our SV dataset to ensure that it was robust to the myriad technical challenges that often plague WGS studies of germline SVs and confound association studies. We observed four main lines of evidence confirming that our SVs were sufficiently high-quality for subsequent association testing. First, the AFs of SVs identified in our study were strongly correlated with the AFs of SVs reported in gnomAD v3.0 (N=63,046 individuals) when matched by ancestry ($R^2$=0.62-0.71; all $P<10^{-10}$; **Fig. S2**) (*8*). Second, the distribution of genotypes for 87-96% of SVs were consistent with Hardy-Weinberg Equilibrium (HWE) within each ancestry. Although there are several *bona fide* biological reasons for why polymorphic variants may deviate from HWE (e.g., recessive or balancing selection), we interpreted this general adherence to HWE as support for the accuracy of our SV genotypes. Third, across 859 complete parent-child trios, 95.8% of all SVs per child were also detected in at least one parent; the remaining 4.2% of SVs per child reflected the sum of proband false positives, parental false negatives, and true *de novo* SVs. Fourth, comparisons to published long-read WGS-based SVs for 24 samples confirmed 90.9% of our short-read WGS-based autosomal deletions after excluding segmental duplications and simple repeats (*32, 33*).

### Supplementary Figures

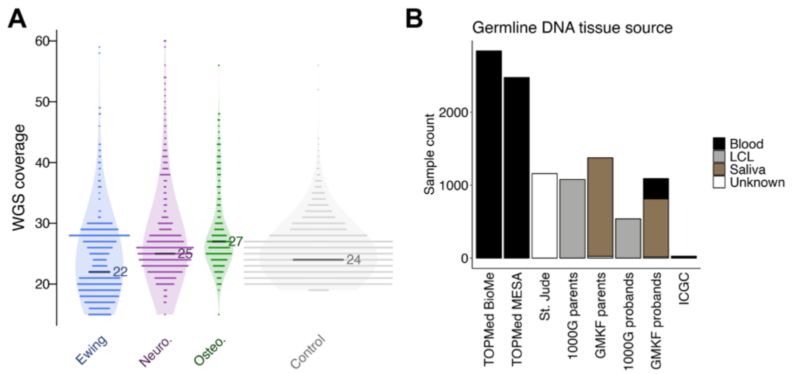


**Fig. S1 | WGS sample properties.** (**A**) Genome-wide coverage for all samples in our final SV dataset stratified by disease. (**B**) Breakdown of germline DNA tissue source for samples across case and control cohorts.

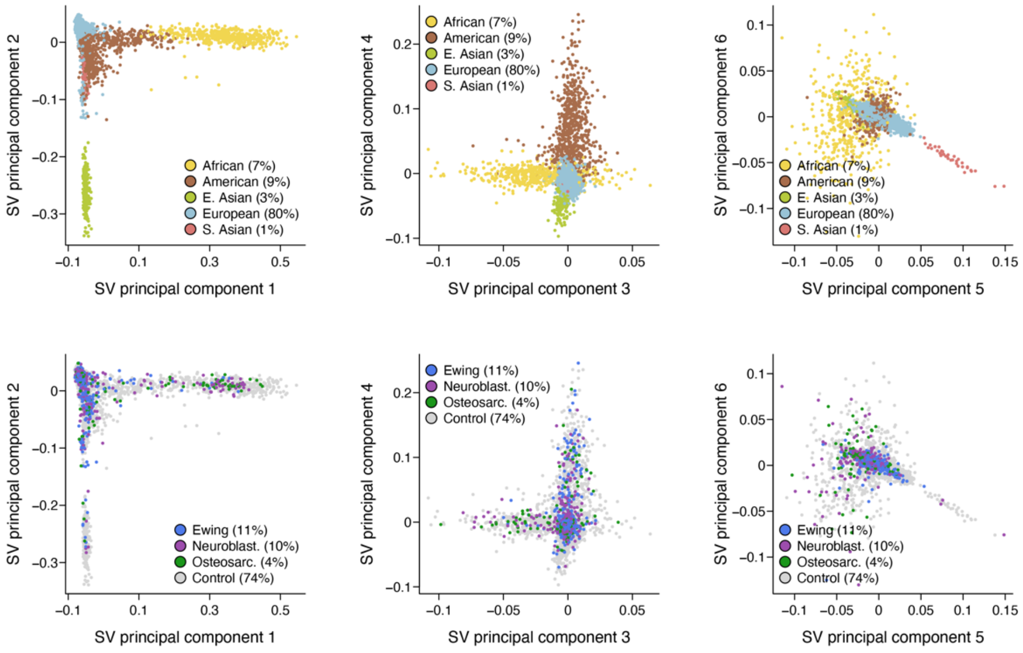


**Fig. S2 | Principal component analysis of common SVs.** We performed PCA on common, well-genotyped SVs in our final SV dataset. Shown here are the top six PCs in the final subset of 6,728 unrelated cases and controls colored by ancestry assignment (top row) and phenotype (bottom row).

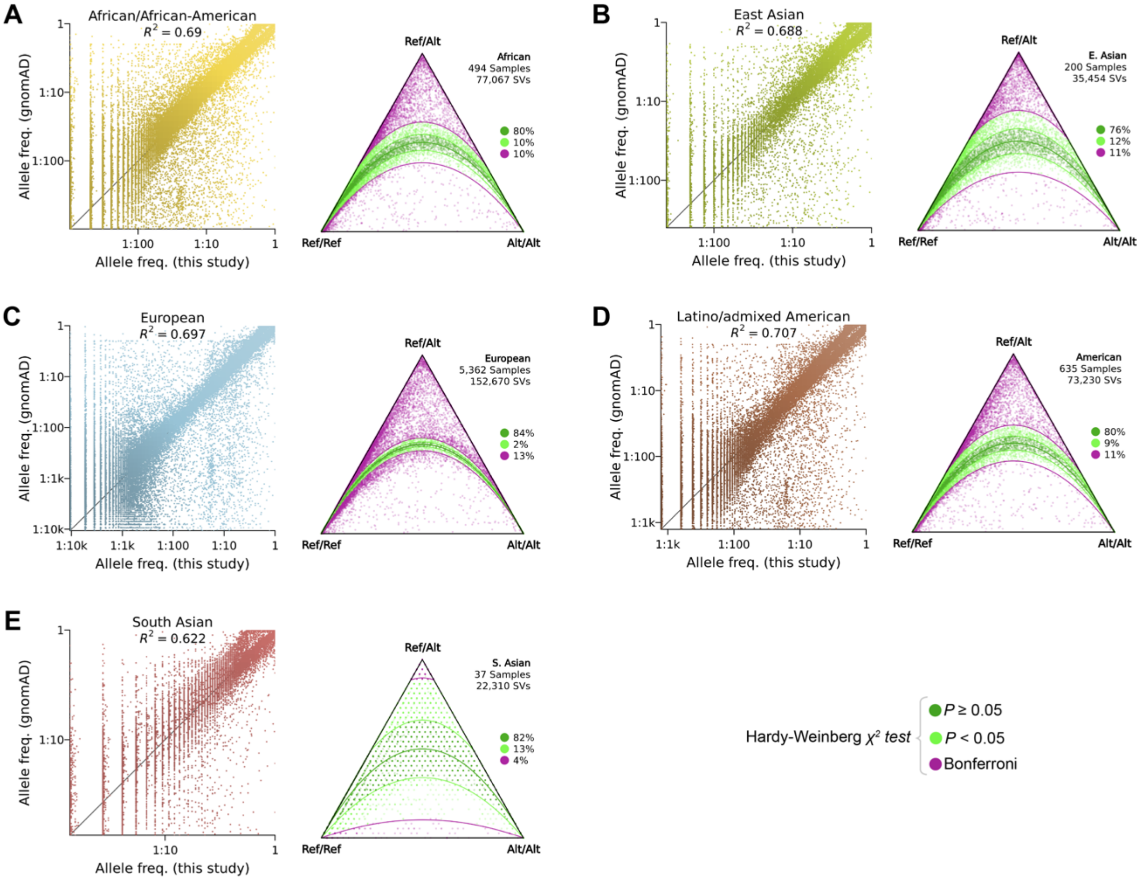


**Fig. S3 | Ancestry-aware SV quality assessments.** We performed two analyses within each continental ancestry group as part of our data quality control and benchmarking (see **Note S1**). Left sub-panels: for each ancestry, we compared the AFs of SVs detected in this study with the AFs of matching SVs reported in gnomAD v3.0 (*8*) and assessed the consistency in AFs between these two datasets with a Pearson correlation analysis. Right sub-panels: we also evaluated the distribution of genotypes for all biallelic SVs present in each ancestry group and tested whether the genotypes were consistent with expectations of Hardy-Weinberg equilibrium (HWE) as a proxy for genotyping accuracy. Each panel depicts these two analyses when restricting our dataset to (**A**) African/African-American, (**B**) east Asian, (**C**) European, (**D**) Latino/admixed American, or (**E**) south Asian samples, respectively.

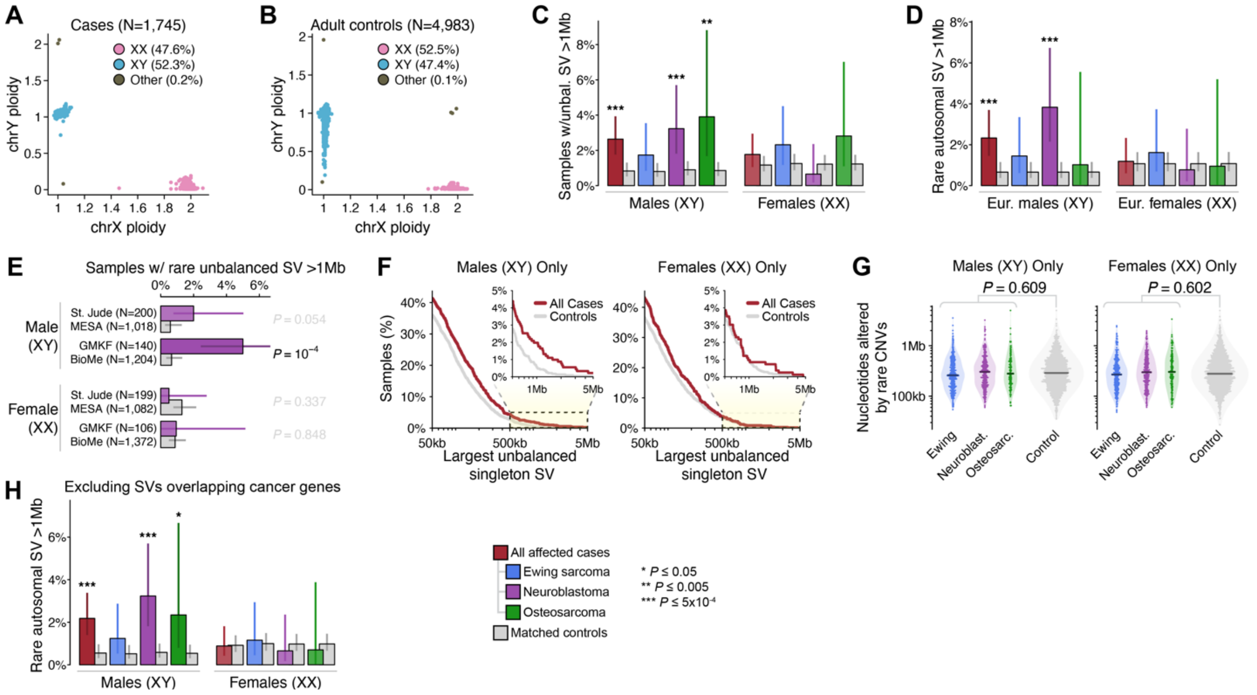


**Fig. S4 | Supporting results for analyses of large, unbalanced SVs.** (**A**) Sex inference among the 1,745 pediatric cancer cases used for formal association testing. (**B**) Sex inference among the 4,983 adult controls used for formal association testing. We observe a clear signature of age-related mosaic loss-of-Y in blood, which is a well-established phenomenon in adult populations (*47*) and is absent in our pediatric cases, as expected. (**C**) Carrier rates by phenotype for all large (>1Mb), rare (AF<1%), unbalanced SVs in males (left; karyotypic XY only) and females (right; karyotypic XX only). See legend at bottom of figure for color key and significance labels. Error bars indicate 95% confidence intervals. (**D**) Carrier rates of large, rare, unbalanced, autosomal SVs in males and females after restricting to individuals of inferred European genetic ancestry. (**E**) Carrier rates for neuroblastoma cases and ancestry-matched controls in males and females when partitioned into independent pairs of case and control cohorts (St. Jude vs. MESA or GMKF vs. BioMe). (**F-G**) Extensions of Fig. 2E-F when separately analyzing males and females. (F) Reverse cumulative distributions of all cases versus controls as a function of the largest unbalanced singleton SV carried in their genomes. (G) Total sum of genomic base pairs impacted by rare, autosomal CNVs of any size per phenotype. (**H**) Carrier rates of rare, large, unbalanced, autosomal SVs after excluding any SV that overlaps a protein coding exon from a gene with an established role in cancer predisposition or progression (*48-51*).

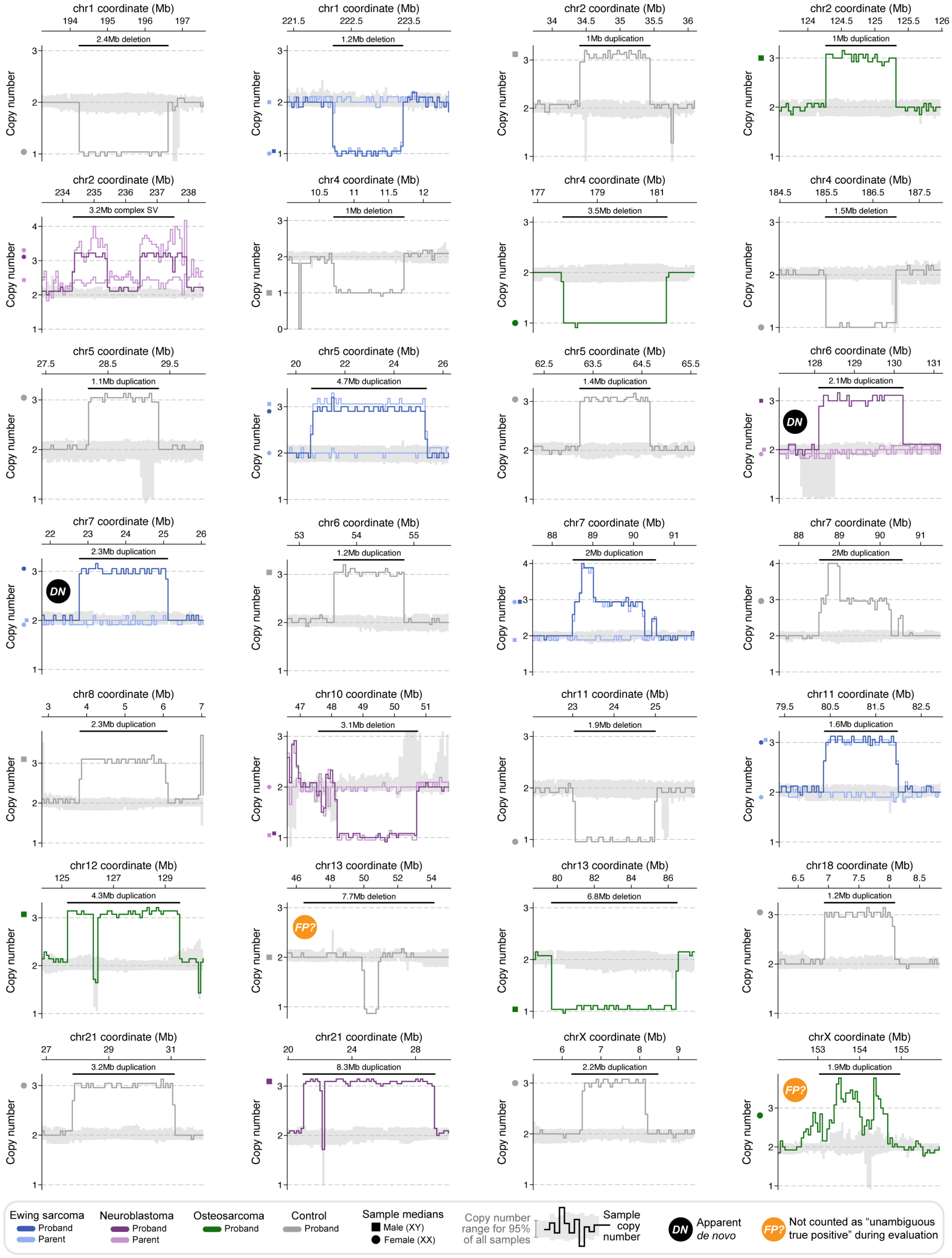
chr1 coordinate (Mb)
2.4Mb deletion
chr1 coordinate (Mb)
1.2Mb deletion
chr2 coordinate (Mb)
1Mb duplication
chr2 coordinate (Mb)
1Mb duplication
chr2 coordinate (Mb)
3.2Mb complex SV
chr4 coordinate (Mb)
1Mb deletion
chr4 coordinate (Mb)
3.5Mb deletion
chr4 coordinate (Mb)
1.5Mb deletion
chr5 coordinate (Mb)
1.1Mb duplication
chr5 coordinate (Mb)
4.7Mb duplication
chr5 coordinate (Mb)
1.4Mb duplication
chr6 coordinate (Mb)
2.1Mb duplication
DN
chr7 coordinate (Mb)
2.3Mb duplication
DN
chr6 coordinate (Mb)
1.2Mb duplication
chr7 coordinate (Mb)
2Mb duplication
chr7 coordinate (Mb)
2Mb duplication
chr8 coordinate (Mb)
2.3Mb duplication
chr10 coordinate (Mb)
3.1Mb deletion
chr11 coordinate (Mb)
1.9Mb deletion
chr11 coordinate (Mb)
1.6Mb duplication
chr12 coordinate (Mb)
4.3Mb duplication
chr13 coordinate (Mb)
7.7Mb deletion
FP?
chr13 coordinate (Mb)
6.8Mb deletion
chr18 coordinate (Mb)
1.2Mb duplication
chr21 coordinate (Mb)
3.2Mb duplication
chr21 coordinate (Mb)
8.3Mb duplication
chrX coordinate (Mb)
2.2Mb duplication
chrX coordinate (Mb)
1.9Mb duplication
FP?
Copy number
Ewing sarcoma
Proband
Parent
Neuroblastoma
Proband
Parent
Osteosarcoma
Proband
Control
Proband
Sample medians
Male (XY)
Female (XX)
Copy number range for 95% of all samples
Sample copy number
DN Apparent *de novo*
FP? Not counted as "unambiguous true positive" during evaluation

(*continued from previous page*) **Fig. S5 | Example read depth profiles for large, rare, unbalanced SVs in pediatric cancer cases and adult controls.** We performed manual review of the RD evidence supporting all 84 rare SVs involving segments of deletion or duplication ≥1Mb that were carried by at least one sample in our subset of 6,728 unrelated, ancestry-matched cases and controls. A representative sample of 28/84 of these SVs are provided here, and were selected mostly at random while ensuring they included (i) an even representation of cases and controls, (ii) both possible false positives (marked with orange tags), and (iii) both predicted *de novo* SVs (marked with black tags). The other 56 SVs are not shown here due to the practical page size limits but all 56/56 exhibited RD evidence qualitatively as strong as any of the apparent true positive SVs shown here (i.e., any panel not marked with an orange tag). Each panel here demonstrates the copy number estimated from WGS RD evidence in 1-3 samples for a locus with a predicted large, unbalanced SV. Dashed horizontal lines correspond to integer copy numbers, which should be expected for true germline SVs. Solid bold lines indicate the copy number in uniform windows across the visualized interval for a predicted SV carrier sample (darker shades) or their parents where available (lighter shades; see legend). Dark grey shaded area indicates the range of copy number estimates for the middle 95% of all samples. Small points on the left axis margin indicate the median copy number estimate for each highlighted sample across the predicted SV interval, which is marked with a bold horizontal black bar.

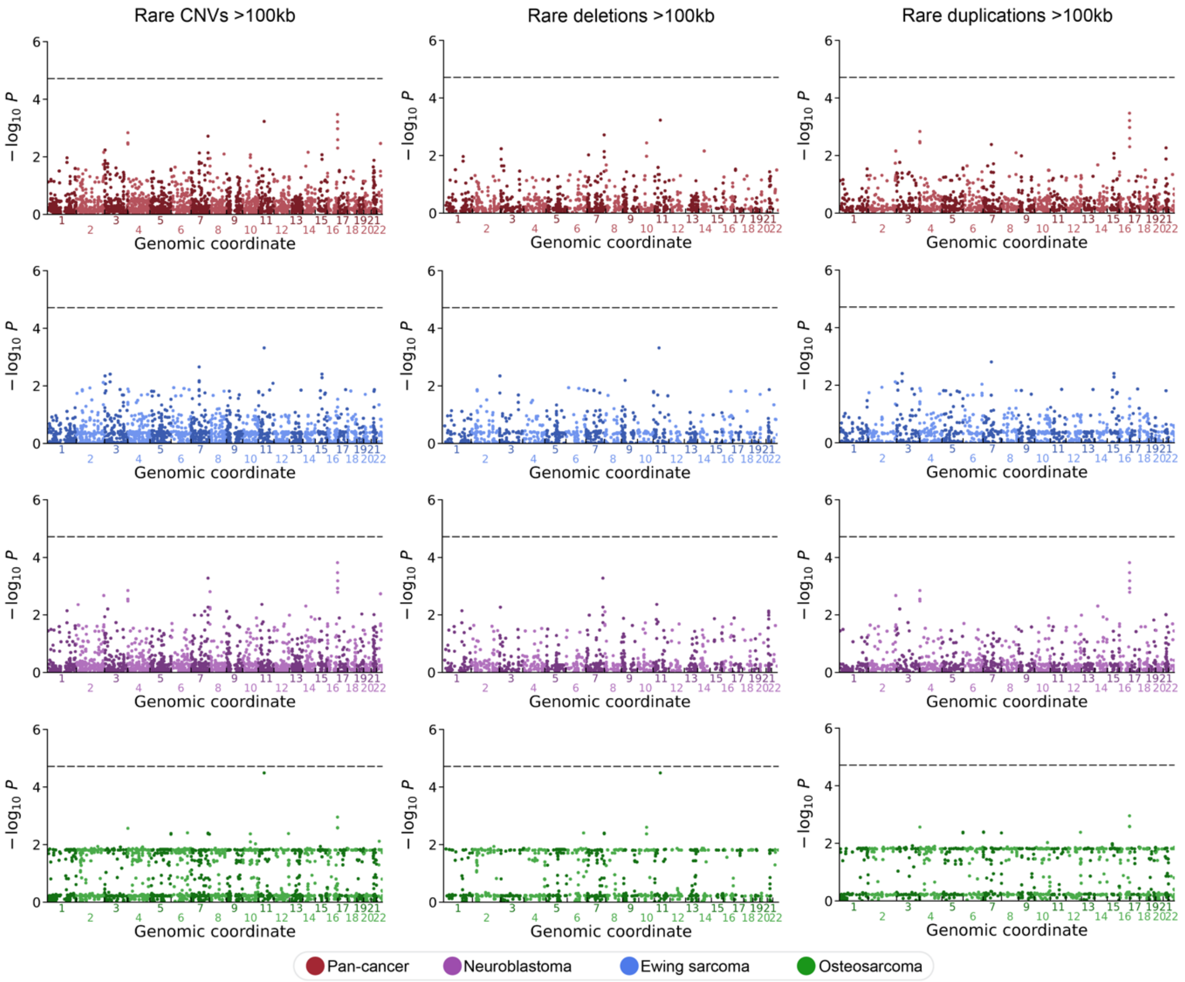


**Fig. S6 | Genome-wide rare CNV association scans.** We conducted genome-wide association scans for each histology (pan-cancer, EWS, NBL, and OS) and rare, large (≥100kb) CNVs using a sliding window approach as previously proposed (*52*) by sliding across all 22 autosomes in 1Mb windows and 250kb steps. For each 1Mb window, we tested for association between rare, large deletions and duplications separately and together for a total of three tests per disease per window. We depict the results for all strata here as Manhattan plots, where each point represents the genomic coordinate and unadjusted P value for a single test. The vertical dashed line reflects a Bonferroni-adjusted P value threshold accounting for all non-overlapping 1Mb windows tested (N=2,623.5 independent windows; $P<1.9x10^{-5}$). No window surpassed this significance threshold for any CNV type in any disease.

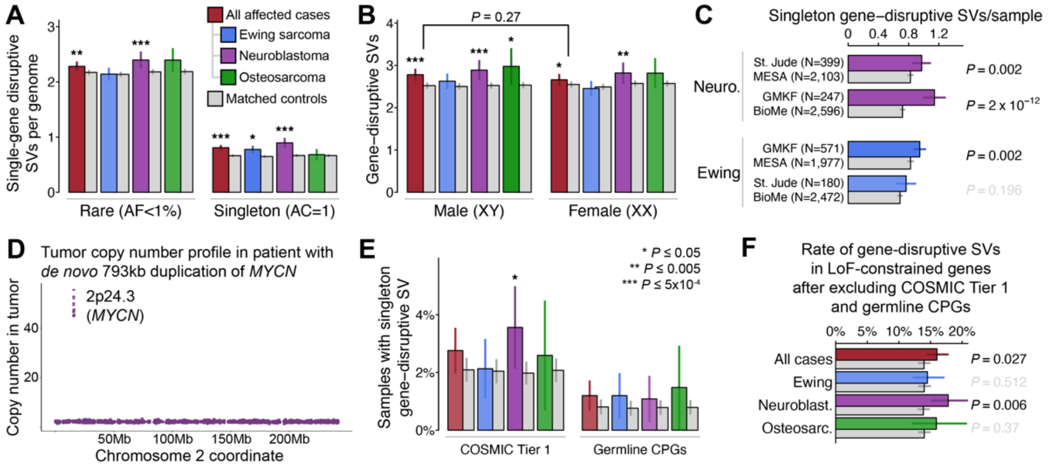

**Fig. S7 | Supporting results for gene-centric analyses.** (**A**) Average number of single-gene-disruptive rare (left) and singleton (right) germline SVs carried in affected cases and ancestry-matched controls. Bars indicate 95% confidence intervals. See (E) for description of P value markers, which were derived from logistic regression adjusted for ancestry, sex, and cohort. (**B**) The enrichment of gene-disruptive SVs per genome did not differ between male and female cases. The P value printed atop the horizontal bracket was calculated using Welch's two-sample t-test. (**C**) The rates of singleton gene-disruptive SVs per genome were consistently higher in neuroblastoma and Ewing sarcoma cases than ancestry-matched controls even when analyzing St. Jude and GMKF cohorts independently. (**D**) We observed a high amplification of *MYCN* in the tumor of the patient with a *de novo* 793kb germline duplication involving *MYCN*. This patient's germline duplication was statistically consistent with a single-copy gain (estimated copy number=3.09; 95% CI=2.78-3.41), which contrasted sharply with the very high copy number amplification (~50 copies) of *MYCN* in the patient's matched tumor; based on these and other data, we concluded that a tumor-normal sample swap was highly unlikely in this case. (**E**) Proportion of samples per disease with singleton gene-disruptive SVs in either COSMIC Tier 1 genes (left) or established germline cancer predisposition genes (CPGs; right) (*48-51*). (**F**) Neuroblastoma cases retained a significant enrichment for rare SVs causing LoF of constrained genes (*38*) even after excluding COSMIC Tier 1 genes and CPGs.

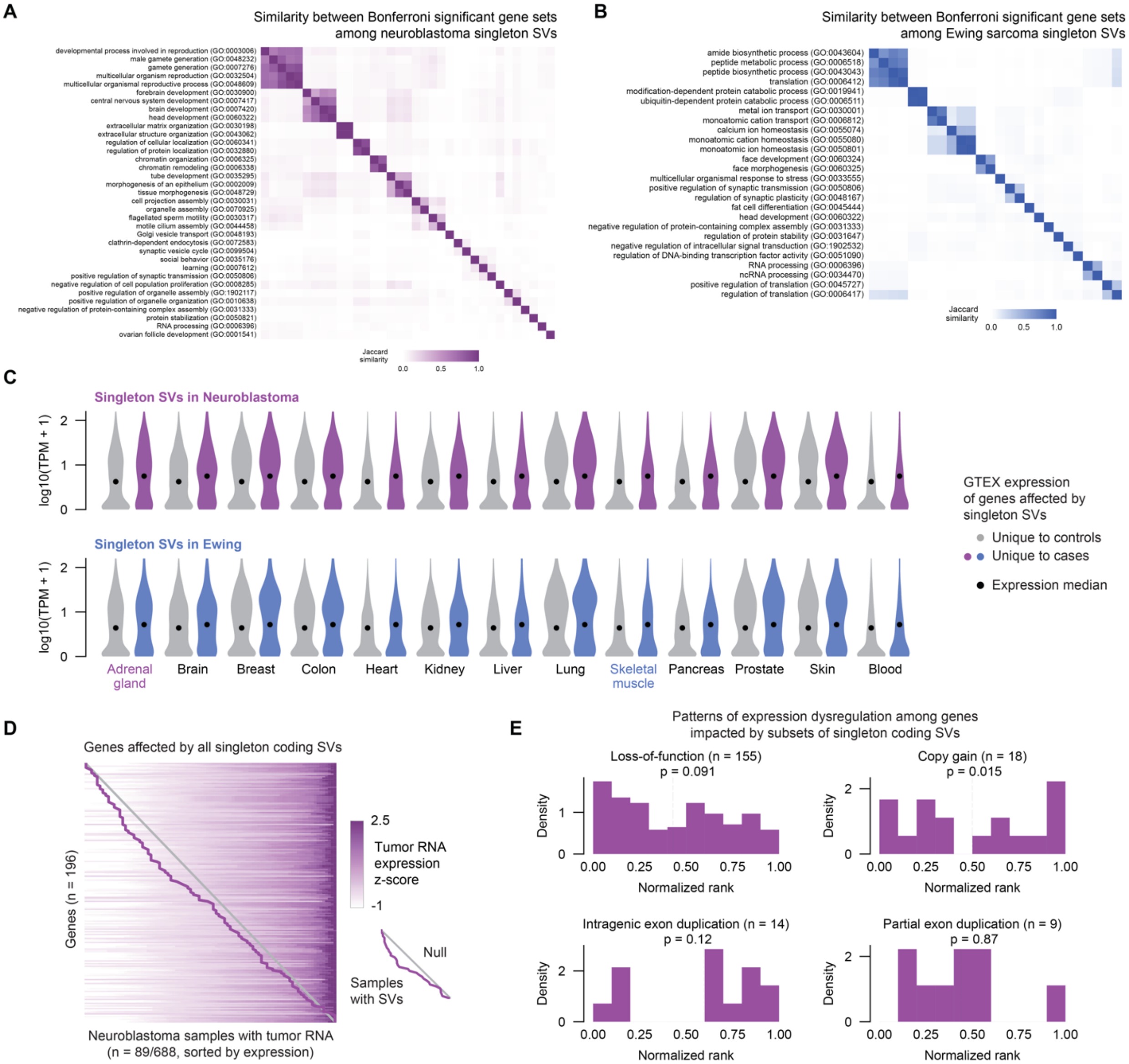


**Fig. S8 | Gene set enrichment and expression dysregulation associated with ultra-rare germline SVs in Ewing sarcoma and neuroblastoma.** (**A-B**) Heatmaps of Jaccard similarity between gene sets that were enriched at Bonferroni significance in coding CWAS of neuroblastoma and Ewing sarcoma. (**C**) GTEx expression of SV-affected genes unique to controls and unique to cases across different tissues in neuroblastoma (top row) and Ewing sarcoma (bottom row). (**D**) Expression heatmap of genes affected by all singleton coding SVs in the subset of neuroblastoma patients with tumor RNA data available. Each row represents a gene and each column represents a sample; samples within each row are ordered by expression z-score. Samples with SVs (one per row) are connected by a purple line. The null uniform distribution is represented by a gray line. (**E**) Individual histograms of all singleton coding SVs, split by predicted effect. Histogram is of normalized rank of SV-affected samples. Black line is mean rank, and p refers to P value from absolute rank deviation.

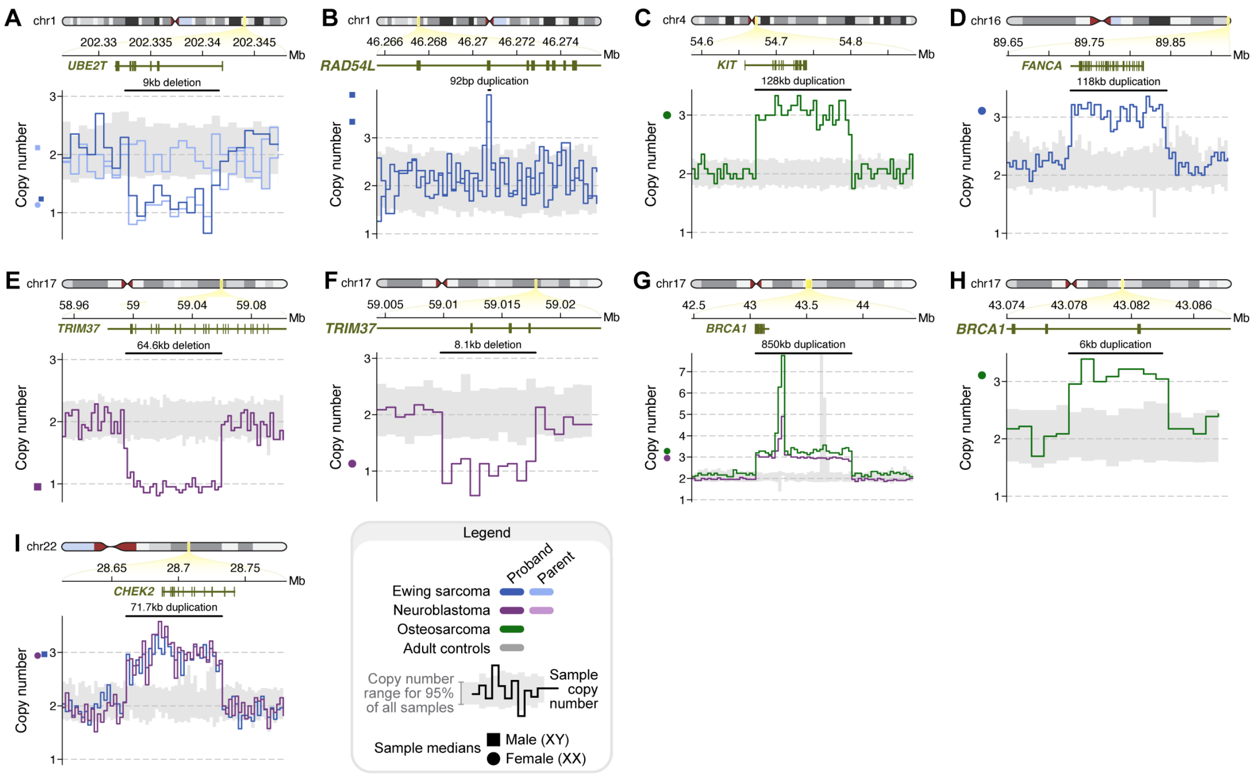


**Fig. S9 | Rare coding germline SVs impacting established cancer genes.** (**A**) LoF deletion of *UBE2T* identified in a child with Ewing sarcoma that was inherited from their unaffected mother. (**B**) Small (92bp) exonic duplication predicted to result in a frameshift of *RAD54L*. (**C**) Partial duplication of *KIT*. (**D**) Whole-gene copy gain duplication of *FANCA*. (**E-F**) An allelic series of two independent rare LoF deletions of *TRIM37* in two unrelated children with neuroblastoma. (**G**) Whole-gene copy gain duplication of *BRCA1*. (**H**) Small, single-exon duplication of *BRCA1*. (**I**) Partial gene duplication of *CHEK2*.

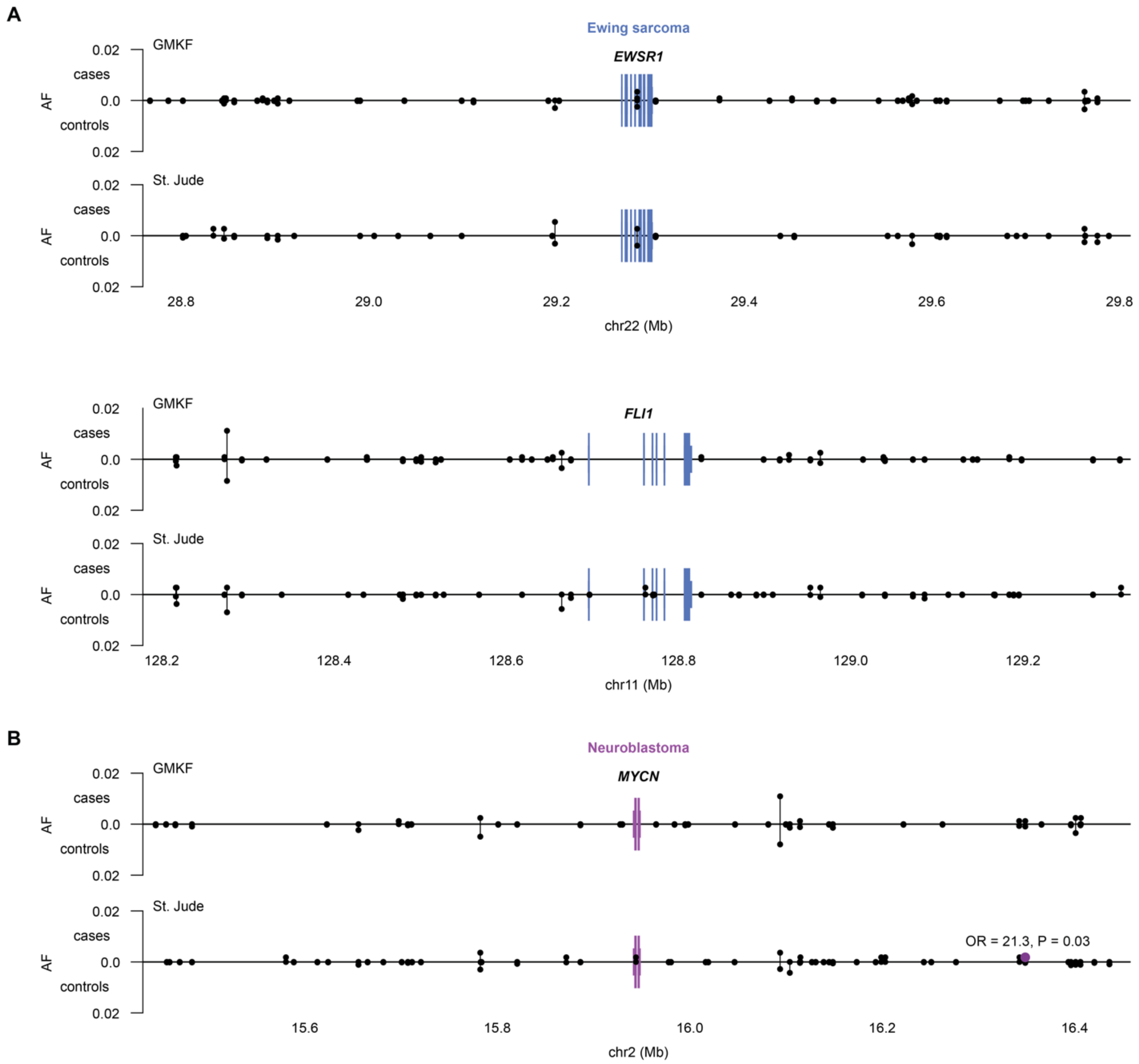


**Fig. S10 | Lack of rare germline SV associations at sites of recurrent somatic SVs in Ewing sarcoma and neuroblastoma.** (**A-B**) Each point corresponds to a single SV, with its frequency in cases and controls encoded by the positive and negative y-axes, respectively. Black points: no significant signal; purple/blue points: nominally significant ($P<0.05$) signal. We observed no associations with rare germline SVs in cases or controls near *EWSR1* or *FLI1* in Ewing sarcoma. There was only a single rare germline SV reaching nominally significant enrichment in cases near *MYCN* in neuroblastoma, which did not remain significant after correcting for the number of SVs tested here.

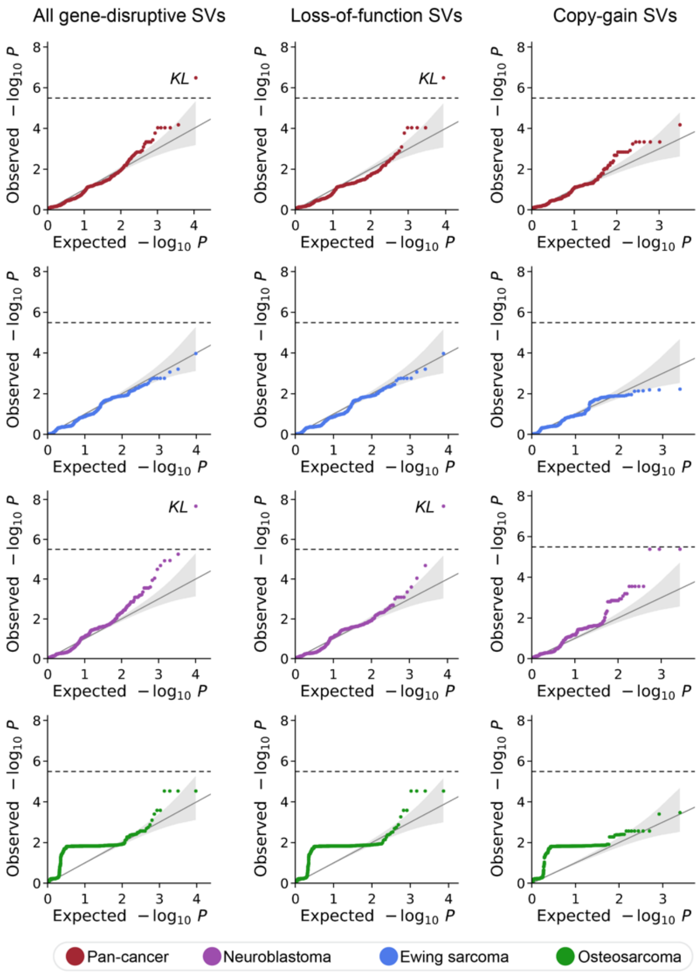


**Fig. S11 | Gene-based coding SV burden tests.** We performed collapsing burden tests for each of 15,544 autosomal, protein-coding genes. One test was conducted per gene for each combination of phenotype (pan-cancer, EWS, NBL, and OS) and predicted SV consequence (all gene-disruptive, LoF, or CG) for a total of 12 tests per gene. The results of these analyses are depicted here as quantile-quantile plots, which compares the distribution of observed P values to expectation under a uniform null. The diagonal line indicates our expectation if there was no true association for any gene, the shaded area indicates the 95% confidence interval of this null expectation, and the dashed horizontal line corresponds to Bonferroni-adjusted P value threshold after correcting for 15,544 genes tested. Just one gene exceeded "exome-wide" Bonferroni significance: rare LoF SVs of *KL* were associated with increased neuroblastoma risk. *KL* is a putative tumor suppressor gene that is silenced in several adult cancers and extends the lifespan of lab mice when overexpressed (*53, 54*).

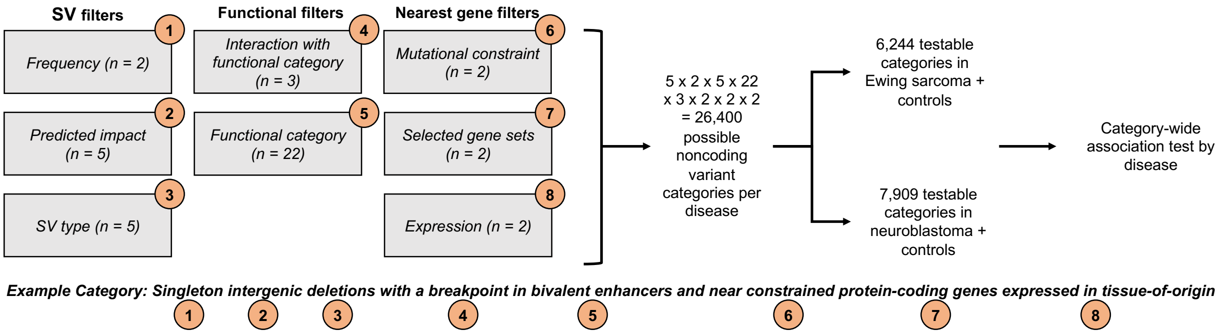


**Fig. S12 | Noncoding CWAS schematic.** We carried out a noncoding category-wide association study (CWAS) in neuroblastoma and Ewing sarcoma (*15, 37*), utilizing eight combinatorial layers of filters to categorize types of noncoding SVs for burden testing. After restricting to categories with >10 SVs observed across cases and/or controls, we retained 6,244 and 7,909 categories of noncoding germline SVs for association testing in Ewing sarcoma and neuroblastoma, respectively.

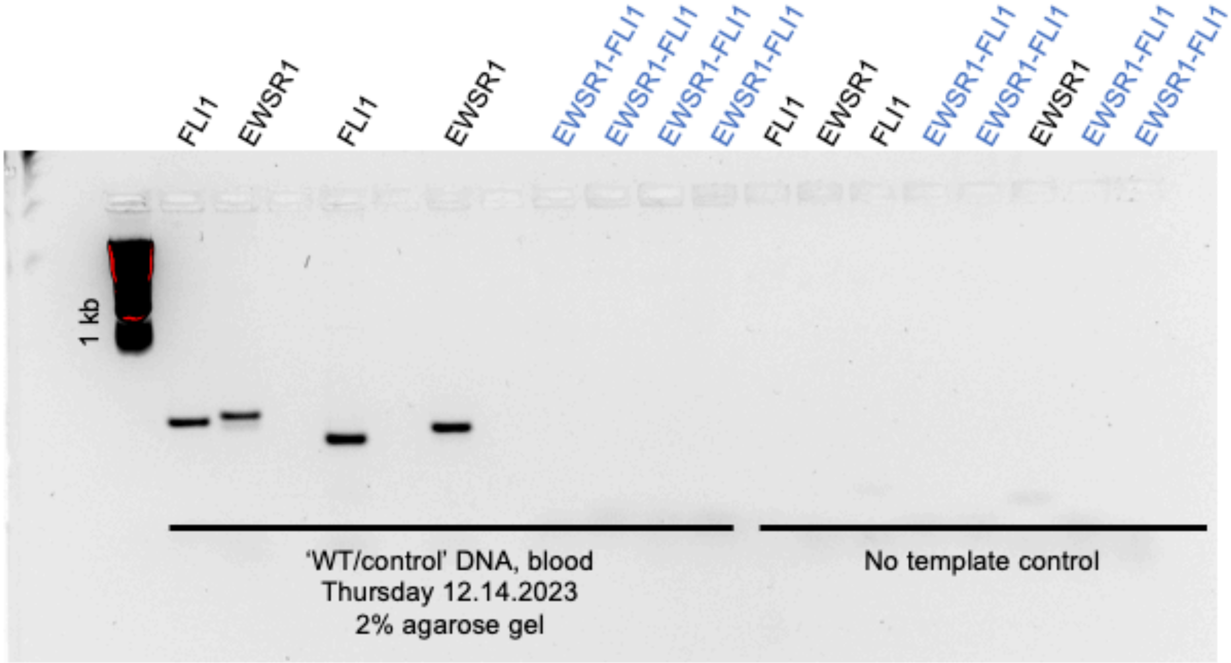


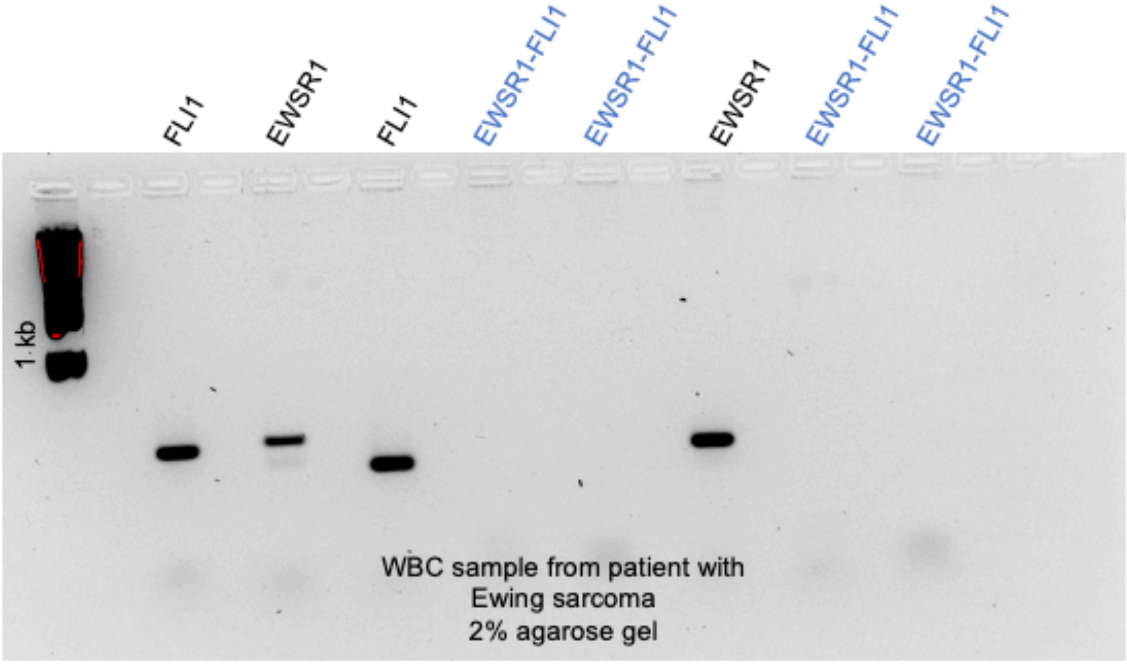


**Fig. S13 | Results of PCR in banked white blood cell DNA from a Ewing sarcoma case with an *EWSR1-FLI1* translocation.** Top: Wild type *EWSR1* and *FLI1* sequences were amplified in a control sample of DNA from an unaffected individual, but not in the non-template control. The *EWSR1-FLI1* translocation sequence was detected in neither control sample. Bottom: In normal tissue from the Ewing sarcoma case, wild type *EWSR1* and *FLI1* sequences were amplified, but the *EWSR1-FLI1* translocation sequence spanning the reported breakpoint was not. This was most consistent with a likely tumor-normal swap and this sample was subsequently excluded from the final germline SV callset.

#### Supplementary Tables

**Table S1 | Cohorts included in study.** List of all WGS datasets included in the present study, including accessions, sample sizes, references, and other information.

**Table S2 | Summary statistics from genome-wide burden tests.** Association summary statistics for selected hypotheses tested in this manuscript. Each row corresponds to a single burden association test for one phenotype.

**Table S3 | CWAS gene lists.** List of protein-coding genes with all annotations (gene sets, constraint, expression) used in CWAS analyses.

**Table S4 | Coding CWAS annotation framework for neuroblastoma.** Combinatorial filters (SV type, SV AF, predicted consequence/genic relationship, genic mutational constraint, putative tissue-of-origin expression, and gene set membership) used for the coding CWAS analysis in neuroblastoma.

**Table S5 | Coding CWAS annotation framework for Ewing sarcoma.** Combinatorial filters (SV type, SV frequency, predicted consequence/genic relationship, genic mutational constraint, putative tissue-of-origin expression, and gene set membership) used for the coding CWAS analysis in Ewing sarcoma.

**Table S6 | Coding CWAS summary statistics for neuroblastoma.** Coding CWAS results for neuroblastoma. Each row corresponds to one category. Effect sizes are captured as log odds-ratios and nominal significance levels are reflected as P values.

**Table S7 | Coding CWAS summary statistics for Ewing sarcoma.** Coding CWAS results for Ewing sarcoma. Each row corresponds to one category. Effect sizes are captured as log odds-ratios and nominal significance levels are reflected as P values.

**Table S8 | Noncoding annotation resources.** Sources of noncoding annotations for the CWAS analyses in Ewing sarcoma and neuroblastoma.

**Table S9 | Noncoding CWAS annotation framework for neuroblastoma.** Combinatorial filters (SV type, SV frequency, SV functional element, SV-element intersection, predicted consequence/genic relationship, mutational constraint of nearest gene, tissue-of-origin expression of nearest gene, and gene set membership of nearest gene) used for the noncoding CWAS analysis in neuroblastoma.

**Table S10 | Noncoding CWAS annotation framework for Ewing sarcoma.** Combinatorial filters (SV type, SV frequency, SV functional element, SV-element intersection, predicted consequence/genic relationship, mutational constraint of nearest gene, tissue-of-origin expression of nearest gene, and gene set membership of nearest gene) used for the noncoding CWAS analysis in Ewing sarcoma.

**Table S11 | Noncoding CWAS summary statistics for neuroblastoma.** Noncoding CWAS results for neuroblastoma. Each row is a category. Effect sizes are captured as log odds-ratios and nominal significance levels are reflected as P values.

**Table S12 | Noncoding CWAS summary statistics for Ewing sarcoma.** Noncoding CWAS results for Ewing sarcoma. Each row is a category. Effect sizes are captured as log odds-ratios and nominal significance levels are reflected as P values.
